# Optimistic and pessimistic cognitive judgement bias modulates the stress response and cancer progression in zebrafish

**DOI:** 10.1101/2024.05.29.596384

**Authors:** Felipe Espigares, M. Victoria Alvarado, Diana Abad-Tortosa, Susana A. M. Varela, Daniel Sobral, Pedro Faísca, Tiago Paixão, Rui F. Oliveira

**Author notes:** These authors contributed equally to this work (Felipe Espigares, M. Victoria Alvarado). **Corresponding author:** Rui F. Oliveira.

## Abstract

Cognitive judgement bias in decision-making under ambiguity occurs both in animals and humans, with some individuals interpreting ambiguous stimulus as positive (optimism) and others as negative (pessimism). We hypothesize that judgement bias is a personality trait and that individuals with a pessimistic bias would be more reactive to stressors and therefore more susceptible to stress-related diseases than optimistic ones. Here, we show that zebrafish judgment bias is a consistent behavioral trait over time, and that pessimistic and optimistic fish express phenotype-specific neurogenomic responses to stress. Furthermore, both phenotypes show differential activation of the hypothalamic-pituitary-interrenal axis in response to chronic stress, suggesting that optimists have a lower stress reactivity. Accordingly, optimists seem to be more resilient to disease than pessimists, as shown by a lower tumorigenesis in a zebrafish melanoma line [Tg(mtifa:HRAS-GFP)]. Together these results indicate that judgement bias is paralleled by differences in the stress response with implications for disease resilience.

## Introduction

Cognitive biases in decision-making have been extensively documented both in animals and humans. A well-studied bias in animal behavior is judgement bias in decision-making under ambiguity, with some individuals interpreting ambiguous stimulus as positive (optimism) and others as negative (pessimism) [1, 2]. There has been a great interest in the study of animal judgment bias under ambiguity to assess animal affective states, based on the principle that negative affective states induce pessimistic bias. Experimental paradigms to assess judgment bias have thus been developed for a broad range of species in recent years [e.g. 3,4], based on the evaluation of an ambiguous stimulus by the animals. These studies have been mainly focused on the modulation of judgment bias by situational or contextual factors (e.g. environmental enrichment) that are expected to affect the affective state of the animals. Judgment bias has therefore been traditionally considered as a transient condition (i.e. a state). However, despite its response to current affective state, it is possible that cognitive judgment bias can also be consistent within individuals (i.e. optimistic vs. pessimistic phenotypes), in which case it may be also considered as a personality trait [5]. Although a number of studies have shown consistency of individual differences in judgement bias across repeated testing [6–8], to the best of our knowledge, no research studies have comprehensively addressed whether judgment bias may reflect a trait, which may be partially due to the conceptual and methodological difficulties inherent to the field of animal personality. However, several tools that were previously developed in the psychological literature and overcome these difficulties have been recapitulated in an integrative theoretical framework [9], which offer a robust and unified approach for the study of personality traits in animals. This framework is mainly based on the analysis of repeatability and several types of the validity (i.e. degree to which a test measures the targeted trait) of the tests that are used to measure the trait of interest.

The hypothesis that judgment bias may reflect a consistent behavioral trait is rather relevant, since it may impact the way individuals respond to and are affected by exposure to chronic stress. In this scenario, it is expected that individuals with a pessimistic bias trait would be more reactive to stressors, because they would consistently interpret ambiguous stimuli as negative, hence over activating the stress response and becoming more susceptible to stress-related diseases than optimistic ones. In fact, patients suffering from stress related disorders often exhibit a pessimistic judgment bias, which frequently lead them to a negative state [10].

Here, we used zebrafish (*Danio rerio*), an emerging model organism in neuroscience and biomedicine [11, 12] to assess the occurrence of consistent between-subject judgement bias, that is the occurrence of optimistic and pessimistic phenotypes in the population, and subsequently to test the hypothesis that individuals with a pessimistic bias trait would be more reactive to stressors and therefore more susceptible to stress-related diseases than optimistic ones. For this purpose, we have first assessed the repeatability and validity of judgement bias to establish that judgment bias is a repeatable behavior that reflects subjective expectations towards ambiguous stimuli. Then, we characterized how optimistic and pessimistic individuals perceive stressors and regulate body homeostasis in response to stress, by documenting their changes in forebrain gene expression and in the hypothalamic-pituitary-interrenal axis, respectively, in response to chronic stress. Finally, we tested the hypothesis that optimists are more resilient to stress-related diseases than pessimists. For this purpose, and based on the evidence that stress can facilitate cancer progression by modulating cancer hallmarks that mediate the transformation of normal cells into malignant cells [13], we have used a zebrafish melanoma line [Tg(mtifa:HRAS-GFP)] to assess the effect of optimism/pessimism on tumorigenesis.

## Results

### Judgement bias is both a behavioral state and a trait

We have phenotyped individual zebrafish for judgement bias using a previously validated test for this species that consists in a Go/No-go task [14, 15]. In brief, fish were trained in a half radial maze to Go to a Positive (P) arm of the maze (rewarded with food) and No-Go to a Negative (N) arm (punished with hand net chase). The P and N arms of the maze were positioned 180° from each other and cued with different color marks (green vs. red), whilst the three ambiguous arms (near positive (NP), ambiguous (A), and near negative (NN)) were positioned at equidistant angles, each separated by 45° between the two reference arms (Fig. 1A). After the training phase the response of fish towards the P and N arms was measured in the absence of the reinforcement cues. After P and N arm testing, the response to the unreinforced ambiguous arm (A) spatially located midway between the two reference arms (N and P; 90°) and cued with a mixed colored card (i.e. half green, half red) was tested. Since an accurate discrimination performance between stimuli (presumably by a generalization response) in the same experimental setup has already been demonstrated in zebrafish [14], a shorter test phase omitting NP and NN cue testing was implemented in this study (Supplemental Note 1). Seventy-three male zebrafish out of the 80 tested (91.25%) learned the discrimination between the P and N arm maze and were subsequently tested for the A arm maze. The response to the A arm shows a bimodal distribution of the frequency of the Judgement Bias Score (JBS), characterized by a lower modal extreme with optimistic-biased fish and a higher modal extreme with pessimistic-biased fish (Fig. 1B). Given this bimodal distribution we have used the lower (optimists) and the upper (pessimists) quartiles of the JBS-frequency distribution for further analyses. The selection of quartiles in this experimental context, as opposed to other divisions such as thirds, offers an efficient data resolution and effectively captures extreme values within our population.

**Fig. 1.**
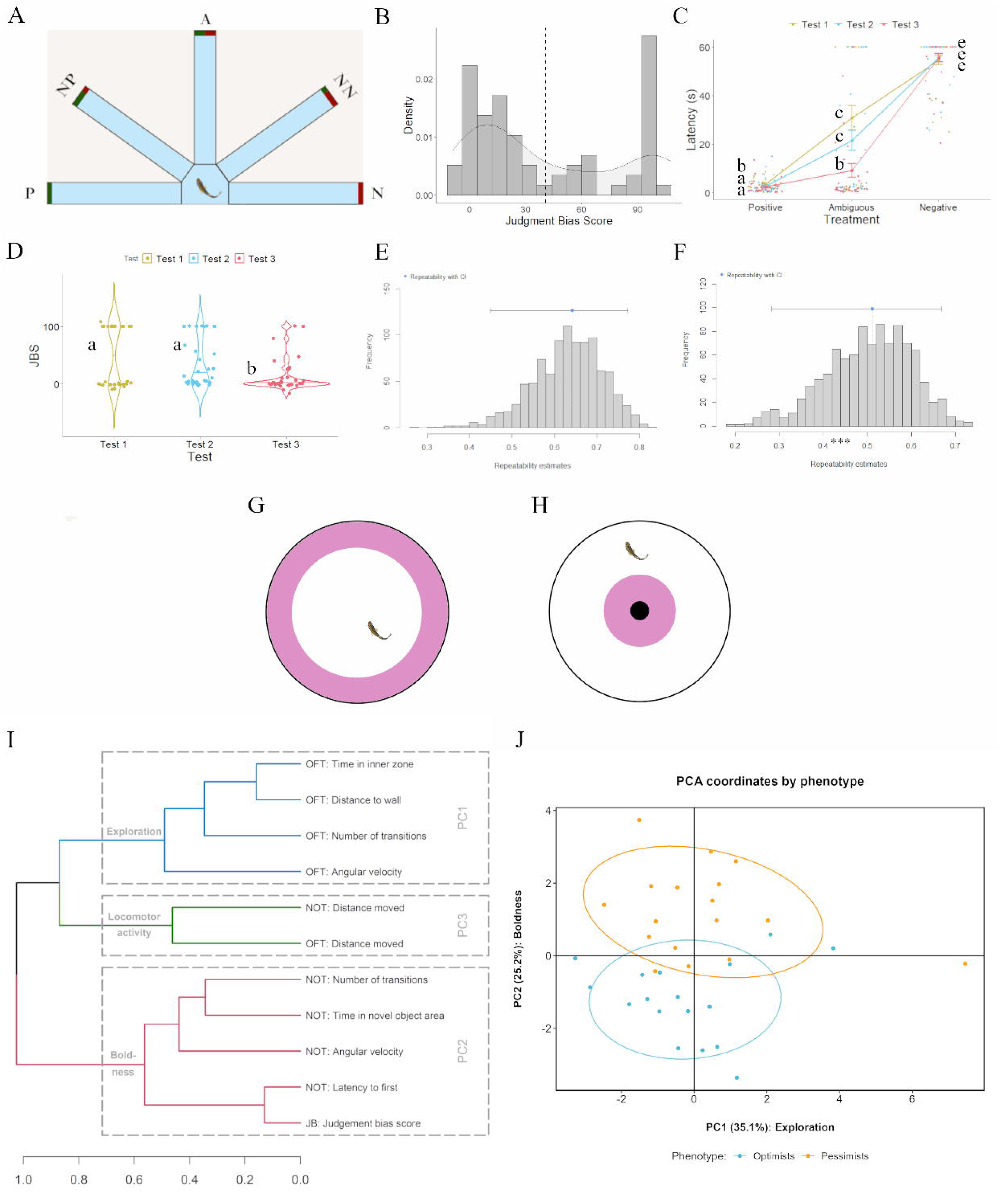
Behavioral characterization of judgement bias in zebrafish. Behavioral characterization of judgment bias in male zebrafish considering the upper and lower quartiles of the Judgment Bias Score (JBS; n = 34). (**A**) Diagram of the half radial maze showing the two reference (positive/rewarded (P) and negative/aversive (N)) and the ambiguous (A) locations that were used in the judgment bias test (JBT) for zebrafish; (**B**) JBS-frequency distribution of the fish that learned the judgment bias task (n = 73); (**C**) Performance of male WT zebrafish in the judgment bias paradigm with repeated testing. Different letters indicate significant differences between the experimental groups (Test 1, Test 2, and Test 3) for each Treatment (P, N, A) following planned comparisons tests. Data are expressed as mean ± s.e.m.; (**D**) JBS of male WT zebrafish with repeated testing. Different letters indicate significant differences between experimental groups (Test 1, Test 2, and Test 3) following planned comparisons tests. Data are expressed as mean ± s.e.m.; (**E**) Bootstrap repeatabilities for the latency to enter the ambiguous arm of the behavioral apparatus; **(F**) Bootstrap repeatabilities for the JBS. (**G**) Diagram of the Open Field Test (OFT); (**H**) Diagram of the Novel Object Test (NOT); (**I**) Degrees of correlation (r) between the different behavioral metrics, illustrated in the cladogram as degrees of association; (**J**) First two components of a Principal Component Analysis (PCA) on all metrics derived from the judgment bias, open field and novel object behavioral tests (JBT, OFT and NOT, respectively), comparing between optimistic and pessimistic fish in a behavioral parameter space. The first two dimensions explain 35.1 and 25.2% of the variance, respectively.

The Judgement Bias Test was repeated over time in order to assess the consistency of the responses to the ambiguous stimulus. Judgement bias has consistently been shown to be modulated by situational or contextual factors that impact an animal’s response to ambiguous stimuli. These external factors include changes in health [16], social dynamics [17], and ageing [14], all of which might inherently compromise trait stability over prolonged periods of time. In fact, a rat study that tested for consistency over 7 weeks obtained a repeatability coefficient for the ambiguous stimulus that although significant, was relatively low (R_ambiguous_= 0.23) [6]. Therefore, to ensure a more comprehensive assessment of trait stability, a three-day testing period was selected (i.e. the test phase was repeated in 3 consecutive days). We believe that shorter durations, as have been established in previous studies [8], allow to capture a robust and reliable representation of the judgement bias trait, minimizing potential confounding effects driven by the above-mentioned external factors. Our results show that the latency to enter the ambiguous arm (A) decreased with repeated testing (i.e. (Test 1 < Test 2 < Test 3; Fig. 1C; Table S1), such that JBS values yielded a decrease in the number of pessimistic responses over time (*p* = 0.005; Fig. 1D; eta^2^ = 0.10, 95% CI = [0.02, 1.00]). It is worth to mention that similar results were found when the whole population (rather than just the upper and lower quartiles) was analyzed (Fig. S1A and S1B Fig; Table S1). Repeatability analysis (i.e. analysis of the intraclass correlation coefficient R) revealed that both measures of judgment bias (i.e. latency to enter the ambiguous arm and JBS) were significantly repeatable when the lower and upper quartiles were considered for the analysis (latency to enter the A arm: R = 0.642; [0.449, 0.773], p < 0.001, Fig. 1E; JBS: R = 0.512, [0.283, 0.67], p < 0.001, Fig. 1F). For the whole population, repeatability of the behavioral measures of judgment bias, was lower but still significant (latency to enter A: R = 0.43, [0.274, 0.568], p < 0.001, Fig. S1C; JBS: R = 0.368; [0.216, 0.499], p < 0.001, Fig. S1D). These data indicates that more extreme judgement bias phenotypes are more consistent. It is worth noting that, as a result of our specific protocol settings, a ceiling effect is observed at the upper extreme of our data, specifically in Test 1. A ceiling effect is more likely to impact extreme values, which suggests that this effect may constrain the variability of JBS within this range and potentially influence the intraclass correlation coefficients. However, due to an apparent decrease in the neophobic response towards the ambiguous stimulus in Test 2 and Test 3 (which evoke a general decrease in the JBS), the ceiling effect becomes residual in such tests, with minimal impact on the intraclass correlation coefficients. In other words, this decrease in neophobic response leads to a broader distribution of the higher JBS scores in Tests 2 and 3, reducing the influence of the ceiling effect on the overall variability of the data and, consequently, on the intraclass correlation coefficients.

The fact that optimism increases with repeated testing for judgement bias indicates that individuals are progressively getting familiar with the A arm, hence decreasing a neophobic response towards it. This result is in line with the common view of judgment bias as a measure of affective state. However, the intra-individual consistency of judgement bias along time, as revealed by the positive correlations of JBS across the 3 days of testing, indicates that it is also a consistent behavioral trait, with individuals keeping their relative expression of the trait within the distribution of the trait in the population.

### Judgement bias clusters with boldness-related behaviors

The individuals screened (n = 73) and selected (n = 17 per experimental group) for judgement bias were also phenotyped for two established animal personality traits: (1) exploratory behavior, using the open field test (OFT) [18] (Fig. 1G); and (2) boldness, using the novel object test (NOT) [19], in which a novel object (a marble) was introduced into the tank (Fig. 1H). Pessimistic fish exhibited higher values for most behavioral measures of exploratory behavior than optimistic fish (Supplemental Note 1, Fig. S2A–S2E, and Table S2). In contrast, optimistic fish approached more frequently and spent more time in the novel object zone (*p* < 0.001 in both cases; Supplemental Note 1, Fig. S2F–2J, and Table S2), suggesting that they have a higher motivation to investigate novel objects (aka boldness) than pessimistic fish.

To assess the phenotypic architecture of judgment bias we performed a principal component analysis (PCA) based on the correlation matrix between the 11 behavioral measures extracted from the 3 above-mentioned behavioral assays (Judgement Bias Test, OFT and NOT). Four principal components (PC) were extracted (cumulative variance explained of 82.7%; Fig. 1I and Table 1): PC1 showed a strong loading of mobility, edge-orienting, spatial preference and spatial examination measured in the OFT, suggesting the occurrence of an exploratory behavior module; PC2 showed a strong loading of JBS as well as of novel object examination measured in the NOT, indicating the occurrence of a boldness behavioral module which includes judgment bias; PC3 showed a robust loading of mobility measured in both OFT and NOT, suggesting the occurrence of a general locomotor/activity module; no additional behavioral modules were identified in PC4. Using the first two components of the PCA to combine all metrics load on PC1 and PC2 in a common behavioral parameter space (Fig. 1J), shows that optimistic individuals are behaviorally differentiable from pessimistic ones. Taken together, these results indicate that the JBS clusters with novel object investigation, thereby establishing a link between the response to ambiguous stimuli and the willingness to take risks.

**Table 1.**
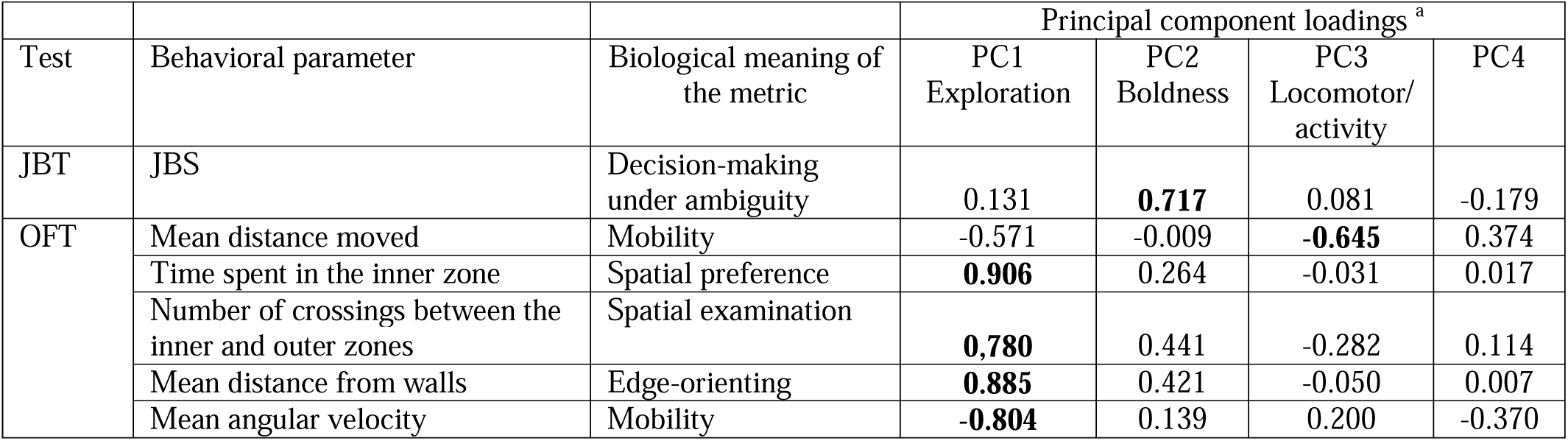
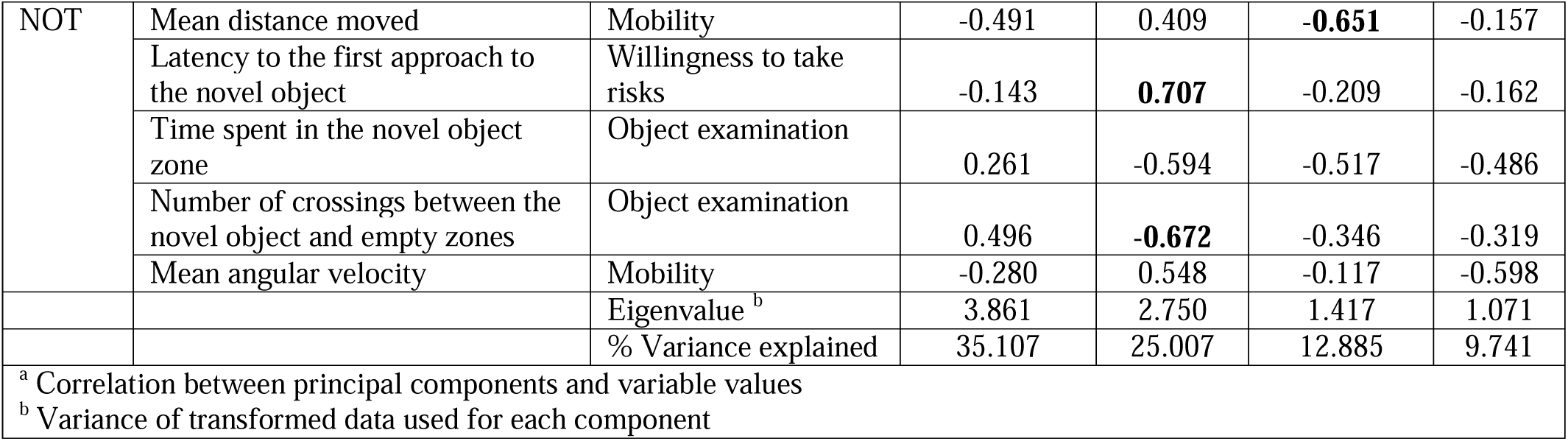
Principal component loadings from the correlation analysis. . Loadings extracted by principal component analysis from the correlation matrix of behaviors across tests for zebrafish belonging to the upper and lower quartiles of the JBS. Bold type indicates the strongest contributors (coefficient > 0.64) to each principal component (PC).

### Genomic brain states of judgment bias phenotypes respond differentially to chronic stress

Given the consistency of judgement bias we tested if the optimistic and pessimistic behavioral phenotypes exhibit different brain states, as measured by the transcriptome profiles of the telencephalon and diencephalon, which are key brain areas for cognitive functioning and physiology and behavior regulation, respectively, and how these brain states respond to stress. Individuals were therefore exposed to a validated unpredictable chronic stress (UCS) protocol for zebrafish during 1 month, and a comprehensive transcriptome analysis, using RNA-seq, of the telencephalon and diencephalon was performed at the end of the experiment using for each experimental treatment (n = 5 fish per experimental treatment: control optimists, control pessimists, stressed optimists, and stressed pessimists stress). The number of animals used in our RNA-seq experiment is consistent with that employed in studies addressing neurogenomic responses in zebrafish [20].

In control conditions there were 21 differentially expressed (DE) genes between optimists and pessimists in the telencephalon and 5 in the diencephalon, whereas after exposure to chronic stress DE genes were 43 and 2, respectively (Fig. S3A–3F; Table S3 and S4). These results suggest a low differential gene expression between the two non-stressed phenotypes, particularly in the diencephalon. In contrast, the transcriptomic response to chronic stress within each phenotype was more pronounced (Fig. 2A–2F). In response to stress optimists have a higher transcriptomic response in the diencephalon (33 and 50 DE genes between control and stressed conditions in the telencephalon and diencephalon, respectively; Fig. 2A and 2C; Table S3 and S4), whereas pessimists have a small response in the diencephalon and a higher response in the telencephalon (48 and 10 DE genes between control and stressed conditions in the telencephalon and diencephalon, respectively; Fig. 2B and 2D). Interestingly, there was almost no overlap in the lists of DE expressed genes in response to stress between optimists and pessimists in either the telencephalon or the diencephalon (Fig. 2E). Looking into the total number of transcripts that were either up-or down-regulated in a phenotype, optimistic fish under stressed conditions had more transcripts down-regulated and fewer transcripts up-regulated in the diencephalon, whereas their pessimistic counterparts had the opposite pattern with fewer transcripts down-regulated and more transcripts up-regulated (Fig. 2F). In the telencephalon, the overall pattern of directional change of pessimists was similar to that of optimists. Taken together, these results indicate that the two judgement bias phenotypes show different brain transcriptomic responses to chronic stress, suggesting differences in cognitive processing of stressors (in the telencephalon) as well as in the regulation of body homeostasis and behavior (in the diencephalon). In fact, the hierarchical clustering of these comparisons (i.e. optimistic control versus optimistic stressed and pessimistic control versus pessimistic stressed; Fig. 2A–2D) indicated that all individuals from each treatment (control vs. stress) were grouped together in separate clusters, reflecting a high consistency of their transcriptome profiles. On the contrary, a consistent clustering was rarely observed in the comparisons between optimists and pessimists either in control or stressed conditions (Fig. S3A–3D). Finally, Gene Ontology (GO) analysis of DE genes detected overrepresentation of specific biological process, molecular processes and cellular component GO terms according to cognitive bias phenotype (Supplemental Note 2, Table S5 and S6).

**Fig. 2.**
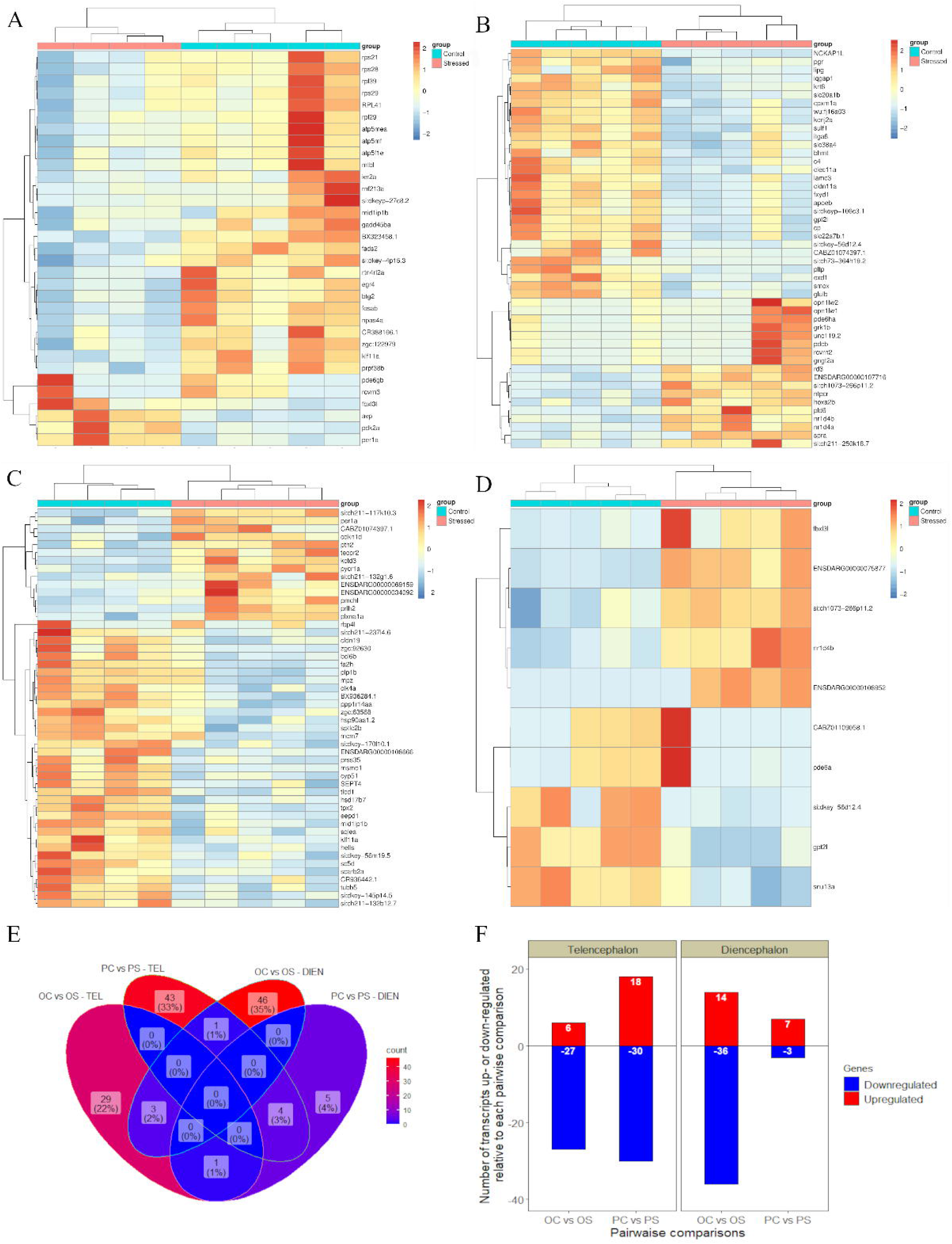
Chronic stress-driven changes in the brain of optimistic and pessimistic zebrafish. Chronic stress-driven changes in gene expression in the brain of judgment bias-related phenotypes (n = 5 per experimental group: optimists control, pessimists control, optimists stress, and pessimists stress). (**A**) Hierarchical clustering of optimistic individuals from each treatment (control versus stress; columns) and of DEG (lines) in the telencephalon; (**B**) Hierarchical clustering of pessimistic individuals from each treatment (control versus stress; columns) and of DEG (lines) in the telencephalon; (**C**) Hierarchical clustering of optimistic individuals from each treatment (control versus stress; columns) and of DEG (lines) in the diencephalon; (**D**) Hierarchical clustering of pessimistic individuals from each treatment (control versus stress; columns) and of DEG (lines) in the diencephalon. Heatmaps represent normalized gene expression levels (red, high expression; blue, low expression); (**E**) Venn diagram showing the number of DEG shared between each control group and their stressed counterparts (control versus stress) in the telencephalon and diencephalon; (**F**) Total number of DEG up-and down-regulated relative to each control group and their stressed counterparts (control versus stress) in the telencephalon and diencephalon.

In summary, the observed divergence in stress-driven changes in neurogenomics states between optimists and pessimists suggests that these two behavioral phenotypes have different responses to stress.

### Judgement bias modulates the response to chronic stress

Given the differences in brain transcriptome responses to stress described above, we tested whether differences in judgment bias may be associated to inter-individual variation in the stress response at different levels of the hypothalamic-pituitary-interrenal (HPI) axis. Therefore, using the same experimental protocol, we also assessed the consequences of chronic stress in optimistic and pessimistic zebrafish at the end of the experiment by measuring cortisol levels under control and stressed conditions as well as expression levels of key genes related to the HPI axis in the brain (i.e. telencephalon and diencephalon), namely corticotropin releasing hormone (*crh*), and glucocorticoid (*gr*) and mineralocorticoid (*mr*) receptors. We also calculated the *mr/gr* ratio, which is a key factor in the regulation of stress reactivity [21].

Chronic stress had a residual main effect (*p* = 0.058; Table S7) on cortisol levels, with stressed fish showing lower cortisol levels than control fish. Indeed, planed comparisons revealed that control optimists and pessimists have similar levels of circulating cortisol, but pessimists decrease their cortisol levels in response to stress (*p* = 0.036) whereas optimists do not (Fig. 3A). Stressed fish, irrespective of phenotype, also displayed higher levels of *crh* transcript levels at both the telencephalon (*p* = 0.052) and diencephalon (*p* = 0.014) (Fig. 3B and 3F). Regarding the expression of *gr* and *mr* receptors in the telencephalon, there was an interaction effect between phenotype (optimists vs. pessimists) and treatment (*p* = 0.008 and *p* = 0.005, respectively). In control conditions, optimists have higher levels of expression of both receptor types than pessimists (*p* = 0.015 and *p* = 0.001, respectively), resulting also in a higher *mr/gr* ratio in optimists (*p* = 0.003) (Fig. 3C–3E). When exposed to stress, optimists decreased *mr* levels (*p* = 0.027), resulting in a significant decrease of their *mr/gr* ratio (*p* = 0.034). In contrast, in response to stress, pessimists showed a significant increase in *gr* levels (*p* = 0.022), and a non-significant trend to increased *mr* levels (*p* = 0.096), resulting in unchanged *mr/gr* ratio in comparison to control conditions. As a result, after exposure to chronic stress there were no significant differences between optimists and pessimists on their expression levels of *gr*, *mr* or the *mr/gr* ratio. Notably, phenotype had a significant main effect (*p* = 0.003) on the *mr/gr* ratio.

**Fig. 3.**
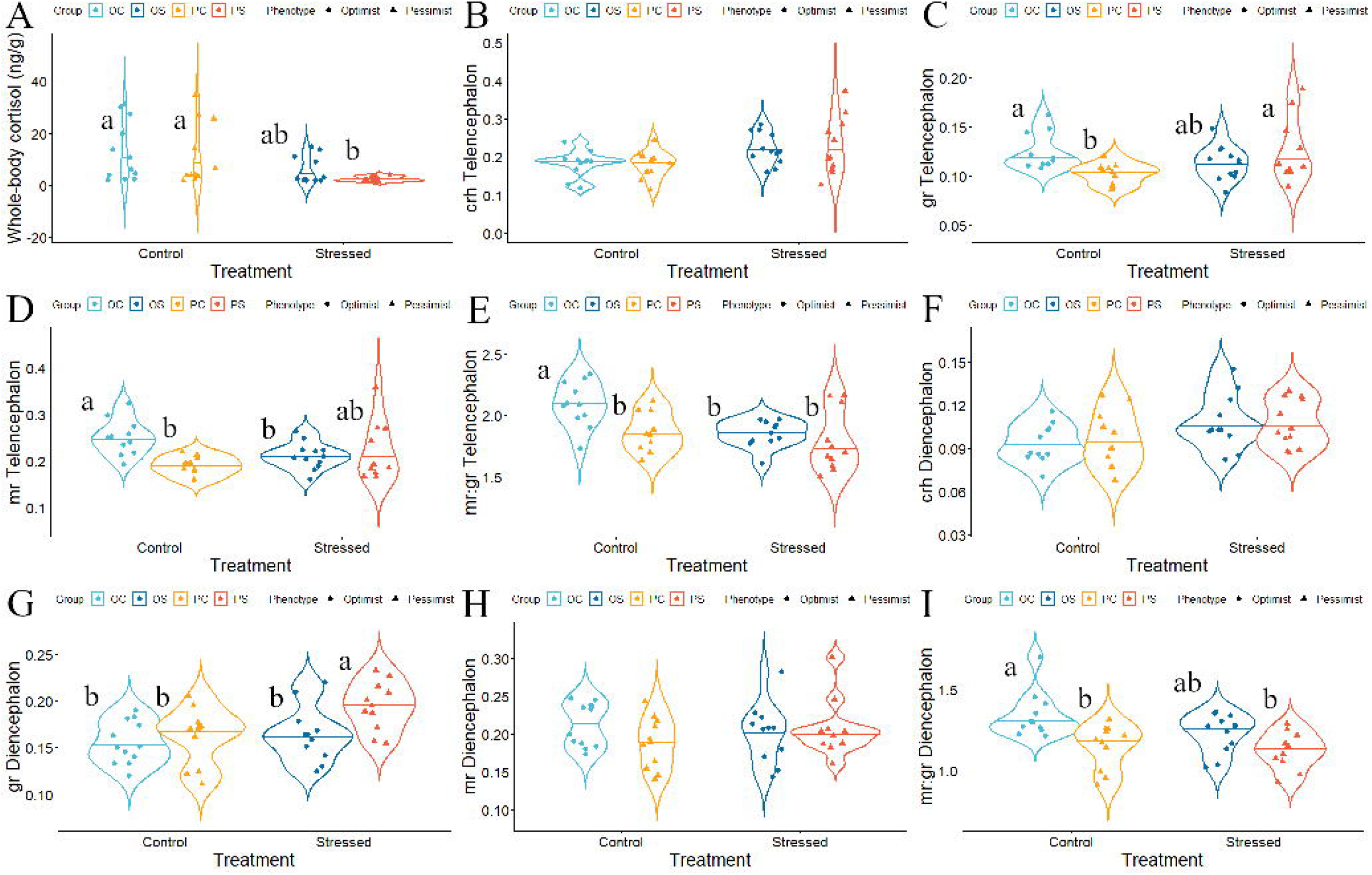
Differential levels of HPI axis-related factors between optimistic and pessimistic zebrafish. Differential levels of a comprehensive set of factors associated with the HPI axis between judgment bias-related phenotypes exposed to the effects of chronic stress (n = 12 per experimental group: optimists control, pessimists control, optimists stress, and pessimists stress). (**A**) Cortisol; (**B**) *crh* expression in the telencephalon; (**C**) *gr* expression in the telencephalon; (**D**) *mr* expression in the telencephalon; (**E**) *mr/gr* ratio in the telencephalon; (**F**) *crh* expression in the diencephalon; (**G**) *gr* expression in the diencephalon; (**H**) *mr* expression in the diencephalon; (**I**) *mr/gr* ratio in the diencephalon. Different letters indicate significant differences between the experimental groups following planned comparisons tests. Data are expressed as mean ± s.e.m.

Regarding the expression of *gr* and *mr* receptors in the diencephalon, there are no significant differences between optimists and pessimists in control conditions, but the *mr/gr* ratio is higher in optimists (*p* = 0.0014) (Fig. 3G–3I; Table S7). When exposed to stress optimists showed no significant changes in either *gr* or *mr* receptor levels, but they showed a non-significant trend for decreased *mr/gr* ratio (*p* = 0.056), whereas pessimists showed increase *gr* levels (*p* = 0.011), but no significant changes in either *mr* or *mr/gr* ratio. As a result, after exposure to chronic stress *gr* levels are significantly lower in optimists than in pessimists (*p* = 0.015), but there were no significant differences between them in either *mr* levels or *mr/gr* ratio. Notably, phenotype had a significant main effect (*p* < 0.001) for the *mr/gr* ratio.

The higher *mr/gr* ratios detected in optimists both in the telencephalon and diencephalon may protect them against chronic stress, hence promoting resilience to disease.

### Judgement bias modulates cancer progression

Given the differential response of optimistic and pessimistic zebrafish to chronic stress, we decided to test if it translates into differential susceptibility/ resilience to disease. Because optimists seem to have a lower stress reactivity, we hypothesized that they should have a lower allostatic load, hence being more resilient to stress-induced disease than pessimists. Based on the known association between stress and cancer we decided to test this hypothesis using a zebrafish melanoma model (mitfa:HRAS) that exhibits full penetrance by 3 months of age [22]. Transgenic (mitfa:HRAS^G12V^-GFP) zebrafish and their WT siblings were phenotyped for judgement bias between 44 and 66 days post hatching (dph), before melanoma onset, using our previously validated protocol [14, 15]. Subsequently, transgenic fish were exposed to chronic stress. Half the optimists and half the pessimists were therefore exposed to the unpredictable chronic stress protocol for 1 month, and the other half of each phenotype remained undisturbed (control group) to assess the effect of stress on tumorigenesis. Individuals were checked weekly for the occurrence of melanoma.

Tg(mitfa:HRAS^G12V^-GFP) individuals (Fig. 4A) are more pessimistic than their WT siblings (Table S8), even before the first signs of melanoma (44-66 dph), as shown by transgenic fish exhibiting increased latencies to enter the ambiguous arm (*p* < 0.001; Fig. 4B) and having higher JBS values (*p* < 0.001; Fig. 4C; Cohen’s d = -0.92, 95% CI = [-1.45, -0.40]). The first tumor incidences were recorded on week 9, which matches previously described tumor onset on this line [23]. Interestingly, in undisturbed fish the cumulative percentage of tumor incidence is significantly lower in optimists than in pessimists at weeks 9 and 10 (*p* = 0.043 and *p* = 0.043, respectively; Fig. 4D), indicating an earlier onset of melanoma in pessimistic individuals (Table S9 and S10). Chronic stress increases cumulative percentage of tumor incidence in optimists (*p* < 0.001) but had no effect on pessimists (*p* = 0.15), which already had a high cumulative percentage in undisturbed conditions, resulting in no difference between optimists and pessimists under chronic stress. At 98 dph all individuals were euthanized and tumor fraction, and PCNA positive tumor cells were quantified. Pessimistic, but not optimistic, individuals showed an increase in tumor fraction in response to chronic stress (Fig. 4E; Table S11), with pessimists showing already a higher level of tumor cell proliferation than optimists, as measured by PCNA, before exposure to stress (Fig. 4F; Table S11).

**Fig. 4.**
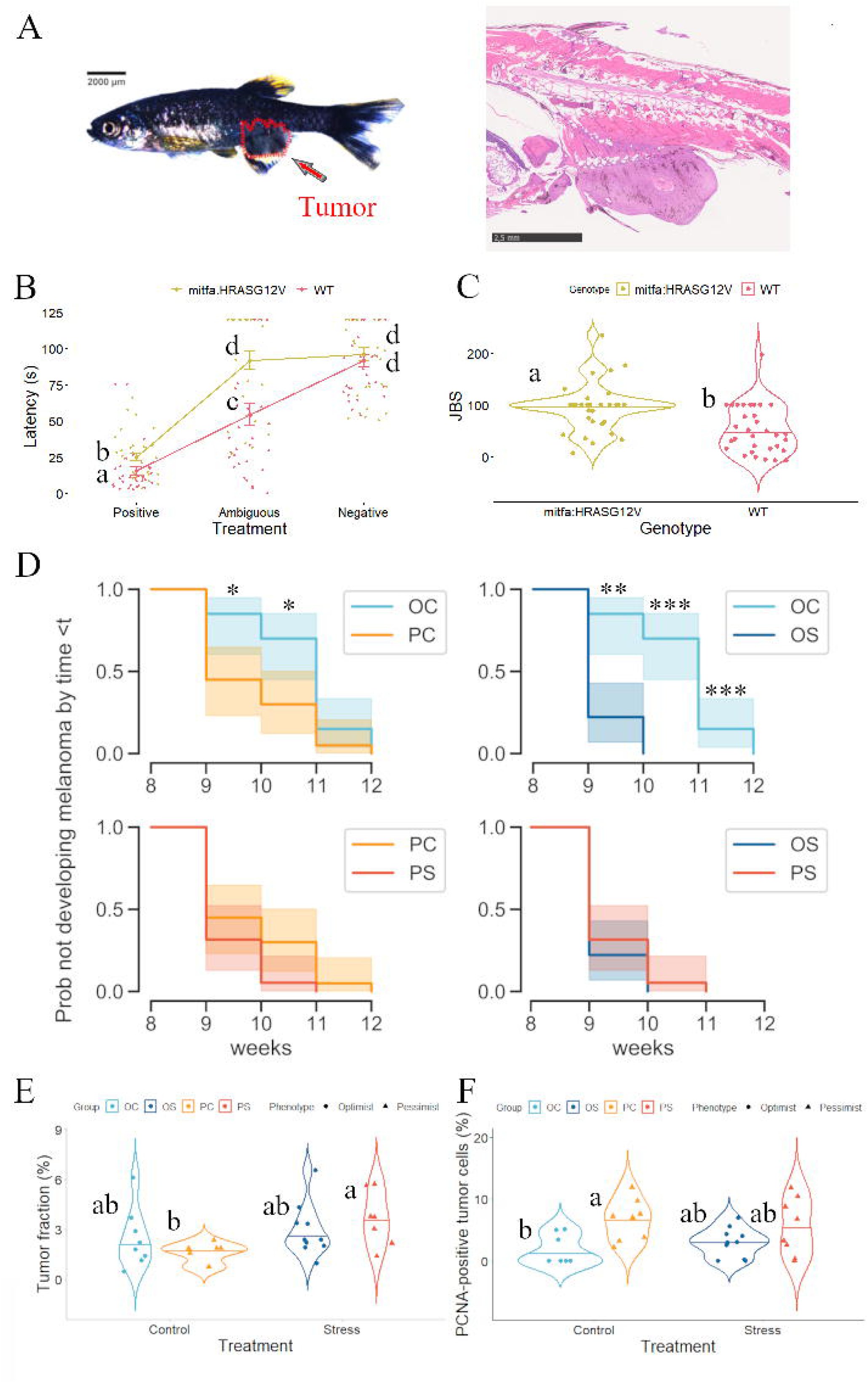
Differences in onset and progression of melanoma tumor between optimistic and pessimistic zebrafish. Differences in onset and progression of melanoma tumor between judgment bias-related phenotypes exposed to the effects of chronic stress (n = 7-11 per experimental group: optimists control, pessimists control, optimists stress, and pessimists stress). (**A**) Representative image of juvenile transgenic zebrafish harboring melanoma tumor and its corresponding histological section; (**B**) Performance of juvenile melanoma zebrafish and their WT siblings in the judgment bias paradigm (n = 32 each group). Different letters indicate significant differences between the experimental groups (transgenic fish and WT) for each Treatment (P, N, A) following planned comparisons tests. Data are expressed as mean ± s.e.m.; (**C**) JBS of juvenile melanoma zebrafish and their WT siblings. Different letters indicate significant differences between experimental groups (transgenic fish and WT); (**D**) Melanoma occurrence over time in juvenile transgenic fish (OC optimists control; PC pessimists control; OS optimists stress; PC pessimists stress). Asterisks indicate statistically significant differences between the experimental groups at each sampling point; (**E**) Tumor fraction (%) in the different experimental groups of juvenile melanoma fish. Different letters indicate significant differences between the experimental groups following planned comparisons tests. (**F**) PCNA-positive tumor cells (%) in the different experimental groups of juvenile melanoma fish. Different letters indicate significant differences between the experimental groups following planned comparisons tests.

The fact that pessimists show a higher susceptibility to tumorigenesis and higher levels of tumor cell proliferation than optimists indicate that optimistic individuals are more resilient to disease than the pessimistic ones.

## Discussion

Our results indicate that judgement bias, usually considered as a behavioral state, can also be seen as a trait (i.e. individually consistent over time), that is correlated with boldness-related behaviors. When exposed to chronic stress pessimistic and optimistic fish presented different neuromolecular responses, as measured by transcriptome changes in the telencephalon and diencephalon, suggesting that they perceive stressors and regulate body homeostasis in response to stress differently. Furthermore, optimists show higher mineralocorticoid receptor/ glucocorticoid receptor ratios in both brain areas, and other differences in the hypothalamic-pituitary-interrenal axis that suggest that they have a lower stress reactivity. Accordingly, optimists seemed to be more resilient to disease than pessimists, as shown by a lower tumorigenesis of optimists in a zebrafish melanoma line [Tg(mtifa:HRAS-GFP)]. Together these results indicate that cognitive bias is a personality trait paralleled by differences in the stress response with implications for disease resilience.

Judgement bias in animals has been seen as a state variable that indicates the affective state of the individuals, and has been widely used in animal welfare studies (i.e. pessimistic bias seen as reporting negative affective state and poor welfare). The occurrence of judgement bias as a trait, that is as a relatively stable behavioral feature of the individual over time (aka personality trait), requires showing significant repeatability across repeated tests over time [24, 25], as well as the establishment of several types of test validity [9]. In this study, we established the repeatability of judgement bias, as well as its ecological validity (i.e. the relationship between the targeted trait and other ecologically important traits). Interestingly, the distribution of JBS in the population shows a bimodal distribution suggesting disruptive selection for the evolution of alternative phenotypes in this judgement bias dimension. Together, these results indicate that judgment bias can be seen as a trait. Furthermore, our PCA analysis indicates that the response to the ambiguous stimulus in the judgement bias test is correlated with the response to the novel object test, positively linking the response to an ambiguous stimulus with the willingness to take risks. A similar link between boldness-related traits and judgment bias has already been reported in a recent study, in which dogs that were rated as more prone to non-social-fear were also more likely to display an optimistic response [26].

The establishment of judgement bias as a personality trait raises the possibility that individuals with different judgement bias phenotypes (i.e. optimists/pessimists) may respond differently to stressors, and therefore express differential resilience to stress-related diseases. We have tested the hypothesis that optimists and pessimists perceive and respond differently to stress by looking into whole-genome changes in gene expression in two key brain regions for the stress response: (1) the telencephalon, which includes brain associative areas that integrate multimodal sensory stimuli and assess the salience and valence of environmental stimuli [27, 28], and (2) the diencephalon, which regulates bodily responses to stress through the modulation of central peptide systems and neuroendocrine axis [29, 30]. Of the differentially expressed genes in the telencephalon of chronically stressed optimists, some of them were identified to have a role in the mediation of the detrimental effects of chronic stress (e.g. *avp*, *fads2*, *klf11a*) and others in sterol metabolism (i.e. *hsd17b7*, *sqlea*, *sc5d*; Gene Ontology Biological Processes ‘Sterol biosynthetic process’ and ‘Cholesterol metabolic process’), which has also been shown to be dysregulated under stress in other organisms [31]. Differentially expressed genes in the brain of stressed pessimists include a set of genes (e.g. upregulated: *nr1d4a*, *nr1d4b*; downregulated: *apoeb*, *kcnj2a*, *cldn11a*) previously linked to stress in multiple species [32–34]. Notably, claudins (*cldn*) have previously been identified as a molecular biomarker of stress with an impact on health and cognition [35]. The fact that chronic stress induces different profiles of gene expression between optimists and pessimists in both brain regions, indicates phenotype-specific neurogenomic responses to stress at the brain level. It is important to note that these data primarily identify neurogenomic differences between these personality traits and stress responses. However, they do not establish a direct link between judgement bias-related phenotypes and stress reactivity, which underscores the need for future research to investigate potential mediating factors involved in stress resilience. In parallel, the neuroendocrine stress axis (i.e. HPI) also showed differences between the two phenotypes. The expression of *crh* in the diencephalon and cortisol levels were used as indicators of the activity of the HPI axis, whereas the expression of the *gr* and *mr* receptors, provided a measure of the sensitivity of the target tissues (telencephalon and diencephalon) to circulating cortisol. In particular, the *mr*/*gr* ratio has been used as a marker of stress vulnerability, since detrimental effects of chronic stress on cognition seem to be GR-mediated, and unbalanced *mr*/*gr* ratios in the limbic brain have been associated to impaired functioning of the emotional and cognitive systems under stress [21]. Higher *mr*/*gr* ratios are therefore indicative of more resilient responses to chronic stress. In baseline conditions there were no differences in cortisol levels between optimists and pessimists, and in response to chronic stress pessimists, but not optimists, decrease cortisol levels. Given that there were no differences between phenotypes neither changes in response to chronic stress in the expression of *crh*, the stress-driven hypocortisolism observed in pessimists does not seem to be due to a down-regulation of the HPI axis but rather to a stress-induced adrenal insufficiency [36]. This interpretation is based on unexpected findings and may benefit from further investigation, while it is true that decreased cortisol levels have been broadly reported in patients with a post-traumatic stress disorder [37] and also in healthy individuals living under stress conditions [38–40]. In baseline conditions, optimistic fish also exhibit a higher *mr/gr* ratio than pessimists, suggesting the latter may have a higher susceptibility to detrimental consequences of chronic stress. Interestingly, under chronic stress, this ratio decreases in optimists but remains relatively stable in pessimists. This intriguing pattern suggests that while optimistic individuals may have an initial advantage with a higher *mr/gr* ratio, they could also harbor a mechanism to attenuate stress-related elevations in glucocorticoids and mineralocorticoids. Therefore, Mr binding in optimists could prevent excessive Gr binding and its associated stress effects, whilst also decreasing *mr/gr* ratio. In contrast, the stability of the *mr/gr* ratio in pessimists under chronic stress may indicate a higher susceptibility to the detrimental consequences of prolonged stress exposure. These results are in agreement with those found in Arctic charr, in which more reactive individuals were characterized by lower *mr*/*gr* ratios in comparison to proactive charr showing decreased tolerance and performance under stress [41].

Given the known link between stress and cancer, we have assessed differences in tumorigenesis in a melanoma line [Tg(mtifa:HRAS-GFP)] between optimists and pessimists exposed to chronic stress. We found a very significant impact of judgement bias phenotype in tumorigenesis in unstressed individuals, with a higher melanoma incidence at an early age in pessimistic individuals. The exposure to chronic stress has a significant impact in tumorigenesis in optimists, that start from a low melanoma incidence, but not in pessimists that start already from high melanoma incidence levels even before stress exposure. Notably, before exposure to stress pessimists already exhibited higher proliferation of tumor cells than optimists, as revealed by PCNA, and after exposure to stress pessimists also shown more enlarged tumors relative to their body size. The link between stress and cancer has been discussed for decades without solid clinical or epidemiological evidence. However, recent animal studies have shown that stress can facilitate cancer progression by regulating molecular and systemic mechanisms that mediate hallmarks of cancer, such as changes in NK cell proliferation, angiogenesis, tumor vascularization and invasiveness and metastasis [42–48]. Moreover, animal studies have shown that exposure to stress during a critical phase of cancer has a greater impact on its progression than non-synchronized stress episodes [13]. This is particularly relevant because in our experiment exposure to chronic stress overlapped with the known melanoma onset period in the zebrafish line used [23]. Our results confirm the role of stress on tumorigenesis and further indicate that between subject differences can be explained by personality traits, such as the judgement bias phenotypes studied here.

In summary, our work highlights the role of psychological factors (i.e. cognitive judgement bias) on the regulation of the brain processing of stress and on the physiological stress response, with an impact in stress-related disease (e.g. cancer). Elucidating how such psychological factors modulate stress vulnerability and its associated disease susceptibility improves our understanding of individual differences in cancer outcomes, and opens the way to conceptualize psychological interventions that might mitigate disease states.

## Methods

### Fish and Housing

Fish used for this study were 4-5 months old male wild-type zebrafish (*Danio rerio*) of the Tuebingen (TU) line and 1.5-3 months old males of the Tg(mitfa:HRAS^G12V^-GFP) line originally developed by the Hurlstone Laboratory (Manchester University, U.K.). All procedures were performed in accordance with the relevant guidelines and regulations for animal experimentation.

### Experiment 1

#### Behavioral assays

Individual zebrafish were first screened in an optimistic – pessimistic dimension by using a previously validated judgment bias test for this species [14, 15, 49]. In brief, a go/no-go task was designed in a half radial maze where individual zebrafish were trained to approach a positive cue (P; food reward) and to avoid a negative cue (N; punishment). Once fish were able to distinguish between P and N cues (as indicated by different latencies to enter each cued arm), their response to an ambiguous (A) cue (an intermediate location/color cue between the P and N locations/color cues) was then tested. In order to assess if judgement bias of the A stimulus of the JBT is associated with other behavioral phenotypes, fish that learned the discrimination between N and P stimuli were selected and tested in an open field test (OFT) and in a novel object test (NOT).

#### Unpredictable Chronic Stress (UCS) protocol

Screened individuals (i.e. optimistic and pessimistic zebrafish) were exposed to a validated UCS protocol for this fish species [50], in which individuals were subjected to a variety of stressors. Each individual experienced two stressors each day, at varying times, for 30 days. All fish of the same experimental tank were given the same stressor at the same time.

#### RNA isolation, qPCR and RNA-Sequencing

Total RNA was isolated from the frozen telencephalon and diencephalon. Isolation of total RNA was performed using the RNeasy® Lipid Tissue Mini Kit (Quiagen), according to the manufacturer’s instructions. The first-strand of cDNA was synthesized using iScript™ cDNA Synthesis Kit (Bio-Rad). All assays were run using 384-well optical plates on a QuantStudio™ 7 Flex Real-Time PCR System (Applied Biosystems™, Thermo Fisher). RNA-Seq libraries were prepared according to the Illumina RNA-Seq protocol and sequenced at the Genomics Unit of Instituto Gulbenkian de Ciência using a NextSeq system to generate single-end 75-bp reads.

#### Cortisol analysis

Whole body cortisol levels were used as a proxy for circulating cortisol levels [51]. Cortisol levels were assessed using an enzyme immunoassay (EIA) kit (Cayman Chemical Company) following the manufacturer’s instructions.

### Experiment 2

#### Behavioral assay and stress protocol

Transgenic zebrafish from 44 to 66 dph (i.e. before melanoma onset [23]) were tested for judgment bias and compared with age-matched WT siblings’ fish. Based on the JBT, transgenic fish that learned the task were classified in an optimistic/pessimistic dimension before entering a UCS experiment (as described in Experiment 1). Fish showing visible tumor masses before entering the judgment bias screening were discarded from the experiment.

#### Tumor assessment

Tumor appearance was assessed macroscopically on a weekly basis, from week 9 (66 dph; experiment start day) to week 13 (98 dph; experiment end date). Individuals were scored for the onset of a vertical growth phase (the lesion develops vertically, forming a nodule), and the presence of an outgrowth in any direction was visually monitored.

#### Tumor measurements

Fish were euthanized the day after the UCS protocol ended. Whole fish were fixed, decalcified, and then processed for paraffin-embedding. Immunohistochemical detection of PCNA was performed using a mouse monoclonal PCNA antibody (PCNA (P10): sc-56; Santa Cruz Biotecnology; dilution 1/100). PCNA-positive cells were revealed by incubation with 3,3’-diaminobenzidine tetrahydrochloride (Liquid DAB+; Palex). Tumor fraction and PCNA positive tumor cells were scored blindly to phenotype and treatment.

### Statistical analyses

The following statistical tests were run in R (v.4.2.2) and SigmaStat (v.3.5). Analysis of the judgment bias testing in WT (Experiment 1) and melanoma fish (Experiment 2) was performed with the R software [52] packages “lme4” [53] and “afex” [54] for the linear mixed effects models (GLMM) and the “emmeans” package [55] for planned comparisons. Generalized linear mixed models (GLMMs) were also used for the analyses of cortisol and stress-related gene expression levels (Experiment 1) and the hallmarks of cancer progression (Experiment 2). A non-parametric two-tailed Kruskal-Wallis test, was used to analyze the JBS in melanoma (Experiment 2) fish. Statistical analyses of the behaviors performed in the OFT and NOT (Experiment 1) were analyzed using a two-way ANOVA followed by an all pairwise multiple comparison procedure (Holm-Sidak’s test).

## Acknowledgments

The authors thank the Fish Facility Platform of the Instituto Gulbenkian de Ciência (IGC) for animal care and the Histopathology Facility of the IGC for technical support of this work.

## Author contributions

Conceptualization: Felipe Espigares, M. Victoria Alvarado, Rui F. Oliveira

Data curation: Felipe Espigares, M. Victoria Alvarado

Formal analysis: Felipe Espigares, M. Victoria Alvarado, Susana A. M. Varela, Daniel Sobral, Tiago Paixão

Methodology: Felipe Espigares, M. Victoria Alvarado

Investigation: Felipe Espigares, M. Victoria Alvarado, Diana Abad-Tortosa, Pedro Faísca

Funding acquisition: Felipe Espigares, Rui F. Oliveira

Supervision: Felipe Espigares, Rui F. Oliveira

Resources: Rui F. Oliveira

Project management: Rui F. Oliveira

Writing—original draft: Felipe Espigares, M. Victoria Alvarado, Rui F. Oliveira

Writing—review and editing: Felipe Espigares, M. Victoria Alvarado, Diana Abad-Tortosa, Susana A.M. Varela, Daniel Sobral, Tiago Paixão, Pedro Faísca, Rui F. Oliveira.

All authors gave final approval for publication and agreed to be held accountable for the work performed therein.

## Data availability

Data is made available as supplementary material to the manuscript.

## Supporting information

### Supplement Methods

#### Supplemental Text

##### Supplementary Figures and Tables

**Fig. S1.** Behavioral characterization of judgment bias in zebrafish considering the whole population (n = 73). (A) Performance of male WT zebrafish in the judgement bias paradigm with repeated testing. Different letters indicate significant differences between experimental groups (Test 1, Test 2, and Test 3) for each Treatment (P, A, N) following planned comparisons tests. Data are expressed as mean ± s.e.m. (B) JBS of male WT zebrafish with repeated testing. Different letters indicate significant differences between experimental groups (Test 1, Test 2, and Test 3) following planned comparisons tests. Data are expressed as mean ± s.e.m. (C) Bootstrap repeatabilities for the latency to enter the ambiguous arm of the behavioral apparatus. (D) Bootstrap repeatabilities for the JBS.

**Fig. S2.** Behavioral differences between fish from the upper and lower quartiles of the Judgment Bias Score (JBS; n = 17 per experimental group: optimists and pessimists): (A) distance moved (cm) in OFT; (B) number of crossings between the inner and outer zones in OFT; (C) time spent in the inner zone (s) in OFT; (D) distance from walls (cm) in OFT; (E) angular velocity (deg/s) in OFT; (F) distance moved (cm) in NOT; (G) latency to first approach to the novel object (s) in NOT; (H) number of approaches to the novel object in NOT; (I) time spent in the novel object zone (s) in NOT; (J) angular velocity (deg/s) in NOT. Asterisks and different letters indicate significant differences between the experimental groups at each sampling point following planned comparisons tests. Data are expressed as mean ± s.e.m.

**Fig. S3.** Phenotypically driven changes in gene expression in the brain of control and stressed zebrafish (n = 5 per experimental group: optimists control, pessimists control, optimists stress, and pessimists stress). (A) Hierarchical clustering of control individuals from each phenotype (optimists versus pessimists; columns) and of DEG (lines) in the telencephalon; (B) Hierarchical clustering of stressed individuals from each phenotype (optimists versus pessimists; columns) and of DEG (lines) in the telencephalon; (C) Hierarchical clustering of control individuals from each phenotype (optimists versus pessimists; columns) and of DEG (lines) in the diencephalon; (D) Hierarchical clustering of pessimistic stressed from each phenotype (optimists versus pessimists; columns) and of DEG (lines) in the diencephalon. Heatmaps represent normalized gene expression levels (red, high expression; blue, low expression); (E) Venn diagram showing the number of DEG shared between each optimistic group and their pessimistic counterparts (optimists versus pessimists) in the telencephalon and diencephalon; (F) Total number of DEG up- and down-regulated relative to each optimistic group and their stressed counterparts (optimists versus pessimists) in the telencephalon and diencephalon.

**Table S1.** Results of the general linear mixed model to assess the effects of Test (Test 1 versus Test 2 versus Test 3), Treatment (Positive versus Ambiguous versus Negative), and the double interaction among these variables. Partial Eta Squared estimates of effect sizes are given for these factors. *Indicates a significant effect.

**Table S2.** Results of the two-way ANOVA to assess the effects of Phenotype (Optimists versus Pessimists), Time (2 min versus 4 min versus 6 min versus 8 min versus 10 min), and the double interaction among these variables. Partial Eta Squared estimates of effect sizes are given for these factors. *Indicates a significant effect.

**Table S3.** List of DEG in the telencephalon.

**Table S4.** List of DEG in the diencephalon.

**Table S5.** Characterization of the DE genes in the telencephalon obtained using ORA for GO, and summarized using GOSlim terms.

**Table S6.** Characterization of the DE genes in the diencephalon obtained using ORA for GO, and summarized using GOSlim terms.

**Table S7.** Results of the general linear mixed model to assess the effects of Phenotype (Optimists versus Pessimists), Treatment (Control versus Stress), and the double interaction among these variables. Partial Eta Squared estimates of effect sizes are given for these factors. *Indicates a significant effect.

**Table S8.** Results of the general linear mixed model to assess the effects of Genotype (mitfa:HRAS^G12V^-GFP versus WT), Treatment (Positive versus Ambiguous versus Negative), and the double interaction among these variables. Partial Eta Squared estimates of effect sizes are given for these factors. *Indicates a significant effect.

**Table S9.** Comparison of the survival curves of the different experimental groups. Treatments were compared using the log-rank test. Correction of p values was performed using the Holm-Sidak method. *Indicates a significant effect.

**Table S10.** Comparison of the survival probability at specific timepoints between treatments that were significantly different. Note that when the survival curve is zero the statistic is infinite. *Indicates a significant effect.

**Table S11.** Results of the general linear mixed model to assess the effects of Phenotype (Optimists versus Pessimists), Treatment (Control versus Stress), and the interaction among these variables. Partial Eta Squared estimates of effect sizes are given for these factors. *Indicates a significant effect.

**Table S12.** Procedure of the unpredictable chronic stress (UCS) protocol in zebrafish.

**Table S13.** Primer sequences, amplicon lengths and annealing parameters for the genes used in the qPCR.

