## Supplementary information for "Optimistic and pessimistic cognitive judgement bias modulates the stress response and cancer progression in zebrafish"

Felipe Espigares *et al.*

**This PDF file includes:**

Supplement Methods

Supplemental Text

Supplemental Figures and Tables

Fig S1 to Fig S3

Table S1 to Table S13

**Supplement Methods**

**Fish and Housing**

Fish used for this study were 4-5 months old male wild-type zebrafish (*Danio rerio*) of the Tuebingen (TU) line (n = 80) (Experiment 1) and 1.5-3 months old males of the Tg(mitfa:HRAS^G12V^-GFP) line originally developed by the Hurlstone Laboratory (Manchester University, U.K.) (n = 216) (Experiment 2). The selection of the sample size for this study was informed by previous investigations conducted within our research laboratory [1, 2]. Drawing upon the collective expertise and knowledge gained from our prior studies in judgement bias, we tailored our sample size to align with the specific objectives and statistical requirements of the current research. Therefore, we believe that our sample size ensures adequate statistical power for robust analyses and meaningful interpretations, while post hoc analyses of effect size further support our findings. Before entering the experiments, individuals were kept in mixed-sex groups (10 adults/L) in a recirculation life support system (Tecniplast^®^) at 28°C, pH 7.0, conductivity of 800 µS/cm, 14L:10D light:dark cycle, and fed twice a day with a combination of live food (*Artemia salina*) and commercial processed dry food (Gemma).

**Experiment 1**

***Judgment Bias Test (JBT)***. Individual zebrafish (n = 80) were screened in an optimistic – pessimistic dimension by using a previously validated judgment bias test for this species that consists in a Go/No-go task [1, 3] (Supplemental Note 4). In order to assess the consistency/reliability of the responses to the ambiguous stimulus (A), the test was repeated 3 times in 3 consecutive days. Fish were video recorded for 1 min during each test trial. Water in the half-radial arm maze was changed between individuals. This behavioral test did not cause any evident impairment of the wellbeing or general condition of the animals.

***Open Field Test (OFT) and Novel Object Test (NOT)***. In order to assess if judgement bias of the A stimulus is associated with other behavioral phenotypes, fish that learned the discrimination between N and P (n = 73) were selected and tested in an open field test (OFT) and in a novel object test (NOT) (n = 17 per experimental group), the day after the JBT was ended (Supplemental Note 4). After testing the fish was returned to its home tank. The water was changed between individuals.

***Behavioral observations****.* Video recordings of the JBT were analyzed using multi-event recorder software (The Observer XT, Noldus technology, version 9). The latency to enter in the target arm was measured for each test trial. A Judgment Bias Score (JBS), which quantifies the degree to which an individual treats the ambiguous cue (A) as the positive or as the negative cue, was computed from the latencies (L) to enter the positive (P), negative (N), and ambiguous (A) arms for each fish as: JBS = (L_A_–L_P_)*100 / (L_N_–L_P_). A JBS over 50 indicates a pessimistic judgment of the ambiguous cue (i.e. fish perceived the ambiguous stimulus as a negative one), while JBS values lower than 50 indicate an optimistic judgement bias (i.e. fish perceived the ambiguous stimulus as a positive one). Given that the JBS standardizes the latency to enter the A arm in relation to individual latencies to enter the P and N arms, it controls for individual motivation, and therefore the possibility that inter-individual differences in the latency to enter the A arm may be linked to differences in swimming speed can be discarded.

Video recordings of the OFT and NOT were analyzed using a computerized video tracking software (Noldus EthoVision XT, Noldus technology, version 12), and several behavioral measures quantified (Supplemental Note 5).

***Individual tagging and experimental design***. Fish with the highest 30 JBS (pessimists) and with the 30 lowest JBS (optimists) were selected for the Unpredictable Chronic Stress (UCS) experiment, while the remaining 13 fish with intermediate JBS were discarded from this experiment. Selected fish were distributed into two groups of five tanks (replicates) each, which received either a UCS protocol (stress group) or were left undisturbed (control group). Each tank consisted of a mixed-phenotype group of 6 fish (i.e. 3 optimists and 3 pessimists). The JBS values were counterbalanced between tanks. A total of four experimental groups were therefore established: optimistic control, pessimistic control, optimistic stressed and pessimistic stressed (n=15 fish per group). Since experimental fish were kept in mixed-phenotype groups throughout the UCS experiment, a common procedure to tag zebrafish with a nylon monofilament was used to facilitate their recognition [4]. Tagged fish were observed daily for 4 days post-tagging to ensure good health and welfare. Fish were then housed in 6-L tanks as previously described.

***Unpredictable Chronic Stress (UCS) protocol***. A validated UCS protocol for zebrafish [5] was used in this study, with small modifications, in which individuals were subjected to a variety of stressors. Each individual experienced two stressors each day, at varying times, for 30 days (Table S12). All fish of the same experimental tank were given the same stressor at the same time.

***Tissue Sampling***. The day after the UCS protocol was completed fish were euthanized with an overdose of buffered tricaine solution (MS-222). Brains were then quickly excised, and the telencephalon (excluding olfactory bulbs) and diencephalon were dissected and quickly snap-frozen in liquid nitrogen and stored at −80°C until use. Whole-body was also collected and kept at −20 °C until use.

***RNA isolation and reverse transcription***. Total RNA was isolated from the frozen telencephalon and diencephalon (n = 12 per experimental group). Samples were lysed in presence of QIAzol Lysis Reagent (Qiagen) and the isolation of total RNA was performed using the RNeasy® Lipid Tissue Mini Kit (Quiagen), according to the manufacturer’s instructions. Purity and concentration of the total RNA was assessed with NanoDrop (Thermo Scientific) and RNA integrity was measured in 10% of samples using a 2100 Bioanalyzer (Agilent Technologies). RNA samples were stored at -80°C until cDNA synthesis. The first-strand of cDNA was synthesized from 0.5 µg of total RNA using iScript™ cDNA Synthesis Kit (Bio-Rad). Samples were then incubated in a thermocycler (T100™ Thermal Cycler, Bio-Rad) according to the supplier’s instructions (5 min at 25°C, 60 min at 42°C and 5 min at 85°C). cDNA samples were stored at -20°C until use.

***Real-time quantitative PCR analysis (qPCR)***. All assays were run using 384-well optical plates on a QuantStudio™ 7 Flex Real-Time PCR System (Applied Biosystems™, Thermo Fisher), using an established reaction protocol (5 min at 95°C, 30 s at 95°C (40 cycles) and 30 s at 72°C). Two technical replicates were performed for each target gene, all run in the same plate. All data were collected and analyzed using the QuantStudio™ Real-Time PCR System software (version 1.1). Elongation factor 1-alpha (*ef1-alpha*), already shown to be suitable for use as a reference gene in zebrafish brain samples [6], was used to normalize the expression of target genes following the equation: 2^CtRef-CtTarget^, where CtRef and CtTarget are the cycle thresholds for the reference gene (*ef1-alpha*) and for the target gene, respectively. Mean of the Cts of two technical replicates were used to calculate the relative expression of each target gene. Two-fold serial dilution of the target genes was used to optimize amplification efficiencies (Table S13), to guarantee an accurate quantification regardless of the DNA template concentration.

***RNA-Sequencing***. RNA-Seq libraries were prepared according to the Illumina RNA-Seq protocol and sequenced at the Genomics Unit of Instituto Gulbenkian de Ciência using a NextSeq system to generate single-end 75-bp reads (20-30 Million reads/sample). Quality of the sequencing data was verified with FASTQC (v0.11.7) [7]. Cutadapt 1.18 [8] was used to remove low quality reads (quality-cutoff set to Q20) and illumine adapter sequences, keeping only paired end-reads having a minimum length of 30 bp. Filtered reads were aligned against the Zebrafish genome (GRCz11) using Hisat22.1.0 [9]. Gene-level counts were generated using Feature Counts 1.6.1 [10] against the Zebrafish gene annotation (Ensembl April 2019). Exploratory data analysis using PCA and sample gene expression correlation matrix identified two samples as outliers (one stressed optimistic telencephalon sample, and one control optimistic diencephalon sample) which were removed from further analysis.

***Cortisol analysis***. Whole body cortisol levels (n = 12 per experimental group) were used as a proxy for circulating cortisol levels [11]. Cortisol levels were assessed using an enzyme immunoassay (EIA) kit (Cayman Chemical Company) following the manufacturer’s instructions and using a 1:4 dilution of the samples. The intra-assay coefficient of variation was 4.30% and inter-assay coefficient of variation was 3.20%.

**Experiment 2**

***Judgement Bias Test (JBT).*** Transgenic zebrafish from 44 to 66 dph (i.e. before melanoma onset [12]) were tested for judgment bias and compared with age-matched WT siblings’ fish (n = 32 each group). Based on the JBT, transgenic fish (from 2 different cohorts) that learned the task (n = 146 out 216) were classified in an optimistic/pessimistic dimension (40 highest JBS = pessimists; 40 lowest JBS = optimists; 66 fish with intermediate JBS were discarded) before entering a UCS experiment. Fish showing visible tumor masses before entering the judgment bias screening were discarded from the experiment (n=27). Since fish from 44 to 66 dph are too small to be individually tagged by the methodology used in Experiment 1, we decided to distribute fish in specific-phenotype groups (i.e. four groups of four tanks each, with 5 optimists or 5 pessimists per tank), which received either a UCS protocol (stress group) or were left undisturbed (control group). This differs from Experiment 1, where heterogeneous phenotypic groups were housed together. While it remains uncertain whether variations in social dynamics between these experimental designs could exist, any potential disparities are not expected to significantly influence the experiment outcomes. This is supported by the fact that chronic stress effects manifest in both experimental settings, with effects consistently aligned, despite differences in social composition. A total of four experimental groups were therefore established: optimistic control, pessimistic control, optimistic stressed and pessimistic stressed (n = 20 fish per group).

***Tumor assessment.*** Tumor appearance was assessed macroscopically on a weekly basis, from week 9 (66 dph; experiment start day) to week 13 (98 dph; experiment end date). Individuals were scored for the onset of a vertical growth phase (the lesion develops vertically, forming a nodule), and the presence of an outgrowth in any direction was visually monitored. Subsequently, individuals were analyzed by histopathology to confirm tumor formation.

***Histological processing.*** Fish were euthanized the day after the UCS protocol ended. Whole fish were fixed in 10% neutral buffered formalin for 72 h at room temperature, decalcified in 0.5-M (ethylenedinitrilo) tetra-acetic acid for 5 days, and then processed for paraffin-embedding. The whole fish was longitudinally sectioned, using a 99 µm interval between sections, and stained for hematoxylin and eosin. Sections immediately following those selected for hematoxylin and eosin staining were used for PCNA immunohistochemistry.

***Whole-slide imaging.*** Whole-slide images were obtained by a NanoZoomer-SQ Digital slide scanner (Hamamatsu Photonics). Measurements of tumor fraction were performed on newCAST stereological software (Visiopharm). Measurement was scored blindly to phenotype and treatment. The results were expressed in volume of fish, volume of tumor, and tumor fraction (volume of tumor/volume of fish). Sex was also determined by histological examination and only males were used given the known sex effect on tumorogenesis and the fact that only males have been used in Experiment 1.

***Measurement of tumor cell proliferation.*** Selected sections were first deparaffinized and rehydrated in order to perform an immunohistochemical detection of PCNA using a mouse monoclonal PCNA antibody (PCNA (P10): sc-56; Santa Cruz Biotecnology; dilution 1/100). PCNA-positive cells were revealed by incubation with 3,3’-diaminobenzidine tetrahydrochloride (Liquid DAB+; Palex). Sections treated with the same protocol but without adding the primary antibody were used as negative control. QuPath software [13] (<https://qupath.github.io>) was used to apply the deconvolution method and the automated cell nucleus detection was performed and optimized visually with a single threshold value for PCNA-positivity (nucleus DAB optic density mean). For each tumor different areas of tumor were selected, which was influenced by the tumor size and the quantity of pigment, but for each tumor, at least 500 cells per tumor were counted, and the measurement was done twice in two different zones. Measurements were scored blindly to phenotype and treatment. The results are expressed as the average of % positive tumor cells.

**Statistical analyses**

Criteria for fish exclusion were pre-established. Fish that failed in the discrimination between P and N testing (as determined by showing a difference < 20 (WT) or < 40 (transgenic fish) seconds in the latency between P and N) were retrospectively excluded for posterior analyses. Furthermore, fish showing an intermediate JBS (between lower and upper extremes of JBS) were discarded for the analysis of gene expression and tumor incidence and progression. Transgenic females were also retrospectively excluded from the experiment after histological determination of sex. Accurate replication was ensured in each experiment. Data were collected in five replicates for each experimental group, which received either a UCS protocol (stress group) or were left undisturbed (control group). Such experimental design was used in both stress reactivity and melanoma experiments. Measurement of tumor cell proliferation was done twice in two different zones. Experimental units were randomly assigned when different treatments were applied. For the judgment bias test, either left or right arm was randomly assigned as the location for positive or negative training, and randomly selected full coloured cards (green or red) were added to these arms. The position (i.e. right or left) and the associated colour cue (i.e. red or green) of the training arms (P and N) were counterbalanced between individuals. For the stress reactivity and melanoma experiments, optimistic and pessimistic zebrafish were randomly distributed between experimental groups (control and stress groups) and experimental tanks (replicates). The JBS values were counterbalanced. Sample sizes were predetermined ahead of time. Specific measures were implemented to mitigate potential bias if applied. Experimenters were not blinded to fish identity during behavioral experiments because animals were run across several behavioral tasks, some of which required long periods of time. Experimenters were not blind to the experimental group assignments of the fish. Experimenters were blinded to treatment conditions for behavioral and gene expression analyses and histological quantification. The following statistical tests were run in R (v.4.2.2) and SigmaStat (v.3.5).

Analysis of the judgment bias testing in WT (Experiment 1) and melanoma fish (Experiment 2) was performed with the R software [14] packages “lme4” [15] and “afex” [16] for the linear mixed effects models (GLMM) and the “emmeans” package [17] for planned comparisons. The response variables were the latencies to respond to stimuli, that is, the time it took the fish to enter the experimental arms [positive (P), negative (N), and ambiguous (A)] of the experimental apparatus. Latencies were restricted to the interval between 0 and 60 seconds (Experiment 1) and 0 and 120 seconds (Experiment 2) and were log-transformed. In Experiment 2, an extension of the test duration was implemented in response to the prolonged latency observed in the transgenic strain when entering all arms of the experimental tank. This adjustment aimed to enhance the capacity for discerning potential variations associated with the different stimuli (i.e. P, N, and A). Importantly, it is worth noting that this modification in test duration is not expected to introduce substantial experimental bias, as latencies were normalized using the JBS. In the model, the fixed effects were Treatment (with three groups: P, N, and A) in interaction with Test (with three groups: Test 1, Test 2, and Test 3) in Experiment 1 or Genotype (with two groups: melanoma line and WT) in Experiment 2. The random effect was the fish identity, since the same individuals were tested in all treatments. A linear model was also used to analyze the JBS in WT fish (Experiment 1). The JBS was log-transformed, and the fixed effect was Test (Test 1, Test 2, and Test 3). Inspection of model residuals showed satisfactory normal distributions. Generalized linear mixed models (GLMMs) were also used for the analyses of cortisol and stress-related gene expression levels (Experiment 1) and the hallmarks of cancer progression (Experiment 2). The response variables were the log transformed concentration of cortisol and the gene expression levels (*crh*, *gr*, *mr* and the ratio *mr/gr*) in the telencephalon and diencephalon. In the telencephalon, *crh* and *mr* expression levels were square-rooted and *gr* expression level was log transformed. Furthermore, tumor fraction was log and % of PCNA positive tumor cells was square rooted transformed. In all models, the fixed effects were Phenotype (Pessimist, Optimist) in interaction with Treatment (Stress, Control). In Experiment 1, random effect was the Tank identity, since the fish of the stressed and control groups were distributed across five tanks (replicates) each. In Experiment 2, random effects were Tank (corresponding to each replicate) and Stock (corresponding to each identical cohort) identities. Inspection of model residuals showed satisfactory normal distributions. A non-parametric two-tailed Kruskal-Wallis test, was used to analyze the JBS in melanoma (Experiment 2) fish. All P-values are two-tailed. Effect size estimates and 95% confidence intervals (CI) were calculated with the R software package “effectsize” [18]. Sample sizes varied either due to technical issues (i.e. sex proportion in experimental groups in Experiment 2) or to the removal of outlier values. The analysis of outliers was conducted for each condition using the generalized extreme studentized deviate procedure with a p = 0.05 and a maximum number of outliers of 20% of sample size. It is important to note that the JBS data was an exception, as no outliers were removed from this dataset. Furthermore, we used principal component analysis for accurate outlier sample detection in our RNA-Seq data [19].

To determine the consistency of the three repeated measurements performed for each focal fish during the JBT (Experiment 1), we performed a repeatability test with the “rptR” package [20]. To estimate repeatability, we employed a linear mixed model using the log transformed latency as the response variable and fish identity as the random effect. To evaluate the uncertainty in the repeatability estimate, we added a bootstrapping argument to the same model with 1000 iterations.

Statistical analyses of the behaviors performed in the OFT and NOT (Experiment 1) were analyzed using a two-way ANOVA followed by an all pairwise multiple comparison procedure (Holm-Sidak’s test). Before the analysis, data transformation was carried out to meet normality and homoscedasticity requirements whenever necessary (OFT distance from walls and NOT distance moved were log transformed; OFT number of crossings, OFT time spent in the inner zone, OFT angular velocity, NOT number of approaches and NOT time spent in the novel object zone were square-rooted transformed; the other variables did not need transformation).

A principal component analysis (PCA) was performed, using the R packages “factoextra” [21], to reduce the number of variables measured in the JBT, NOT and OFT tests (Experiment 1) to a set of principal components (PCs) that represent linear combinations of the original variables, conserving the maximal data variation. The “ggbiplot” package [22] was used to graphically represent individual scores over the first two PCs, categorized by their JB phenotype. A cluster analysis was then performed with the “philentropy” package [23]. We used the Euclidean distance to compute the matrix of distances between all PCs. Then, we applied the hierarchical clustering with complete-linkage. Finally, to plot the dendrogram, we used the “dendextend” package [24].

Differential gene expression analysis (Experiment 1) was performed using the edgeR package [25] within the R software. First, genes with low expression (less than one count per million in any sample) were filtered out. Gene counts of expressed genes were normalized using the trimmed mean of M-values (TMM) normalization method. Differentially expressed genes between the different conditions were obtained using a generalized linear model (GLM) likelihood ratio (LR) test. P-values were adjusted for multiple testing using the Benjamini and Hochberg method. Genes with adjusted p-value of less than 0.05 were considered as differentially expressed. Zebrafish gene ontology (GO) annotation was obtained from Ensembl Biomart. GO term enrichment was performed comparing the GO terms annotated for genes detected as differentially expressed against the full annotation of expressed genes, using a hypergeometric test (the main GO hierarchies were considered separately, and GO terms with one gene – singletons – were not considered).

We performed survival analysis of tumor incidence between experimental groups (Experiment 2) using the log-rank test. The estimated *p*-values were then corrected by the Holm–Sidak method, to account for multiple comparisons and control Type I error inflation.

**Supplemental Text**

Supplemental Note 1

In this study, we implemented a shorter test phase in which NP and NN cue testing was omitted. A shorter test phase and, consequently, a lower number of training trials in this phase could have several advantages in terms of minimizing potential events affecting the categorization of the ambiguous cue. For instance, a higher number of positive outcomes (i.e. food rewards) may lead to a decrease in appetite, which could affect the performance of optimistic behaviors independently of the affective state. Appetite impact on judgement bias tasks has been already reported [26, 27]. On the other hand, a higher confounding influence of stress could be achieved by increasing the number of negative outcomes (i.e. punishments) and/or the overall duration of the test phase. The effects of stress on task learning in judgement bias tests have also been previously reported [28, 29].

Supplemental Note 2

We tested individuals in the lower and upper quartiles of the JBS distribution (n = 17 per experimental group) in the Open field test (OFT), which measures activity/locomotion and exploration in individuals exposed to a novel environment. We examined several measures of activity and location during the whole OFT trial (10 min). No statistical differences were observed in the distance travelled of optimists versus pessimists when tested in this assay (Fig. S2A; Table S2). Phenotype (optimists vs. pessimists) had a significant main effect for the number of crossings, time spent in the inner zone and distance from walls (*p* < 0.001, *p* = 0.002 and *p* < 0.001, respectively), with pessimistic fish exhibiting higher values of these behavioral parameters than optimistic fish. Planned comparisons show that, upon introduction into the novel environment, both phenotypes displayed a predisposition to remain near the center of the arena as well as to perform a high number of crossings between the outer and the inner zones (Fig. S2B-D). Over time, fish gradually swam towards the walls and decreased the number of crosses between zones. Interestingly, pessimistic fish showed significant higher values of these behavioral parameters in the early phase of the assay, coinciding with the period in which both phenotypes exhibited a high propensity to explore the novel environment. Similar to the distance travelled, no difference was observed between phenotypes in angular velocity (Fig. S2E). These results suggest that pessimistic fish exhibit a higher propensity to explore novel environments than optimists.

Subsequently individuals were tested in a novel object test (NOT), which is a standard paradigm used to assess boldness in mammals and fish, in which a novel object (in our study a marble) is introduced into a familiar environment. In our experiment, no statistically significant differences were found in the distance travelled between optimistic and pessimistic fish (n = 17 per experimental group) during the whole NOT (10 min; Fig. S2F). Optimistic fish showed a lower latency to approach the novel object (*p* = 0.012), approached it a higher number of times, and spent a higher proportion of time next to it in the first 4 - 6 minutes of the trial, than pessimists (Fig. S2G-I). In fact, phenotype had a significant main effect for the number of approaches and time spent in the novel object zone (*p* < 0.001 in both cases). Interestingly, pessimistic fish exhibited significant higher angular velocities during this time-bin (Fig. S2J). These results suggest that optimistic fish have a higher motivation to investigate novel objects (aka boldness) than pessimistic fish.

Supplemental Note 3

Gene Ontology (GO) analysis of DE genes in the telencephalon (Table S5) detected a biological process enriched in only optimistic individuals that is related to peptide biosynthesis and protein metabolism. Molecular processes were also only overrepresented in optimists and included terms related to structural integrity. Regarding cellular component GO terms, only optimists had an overrepresentation of DE genes in the ribosome, rough endoplasmic reticulum and cytosol. (GO) analysis of DE genes in the diencephalon detected several biological processes enriched only in optimistic individuals, which are related to sterol and alcohol metabolic processes, and response to stimulus (Table S6).

Supplemental Note 4

Detailed description of behavioral tests:

Judgement Bias Test **-** The judgment bias test (JBT), used a half radial maze (Fig. 1A) where, after a period of acclimatization to the apparatus, fish were trained in the two references arms of the maze to perform different responses when a specific cue was presented (location within the maze arm/color cue). The two reference arms, positive (P) and negative (N), were positioned 180° from each other, while colored cards (i.e. green and red) were associated with each of these arms. During the training phase, individuals learned therefore to perform a Go response towards the P arm when a specific cue (location/color cue) was presented in order to experience a positive event (i.e. food reward), whereas a No-go response towards the N arm should be performed in association with the other cue (location/color cue) in order to avoid a negative event (i.e. punishment by capturing them with a hand net and gently shaking them inside the net for 10 sec). The first session of the Training phase was performed on day 1 after the Habituation phase (inter-phase interval of 4-6 hours), while the second session of such phase was performed on day 2. Each training session consisted of eight entries in total (four negative (N) and four positive (P)) in a pseudo-random sequence (i.e. P P N N P N P N). The Test phase was performed on day 3 and consisted of 6 pre-training trials (*i.e*. P P N N P N) followed by the presentation of P and N cue tests. During this phase, the response of fish towards the P and N arms was measured in the absence of the reinforcement cues. After P and N arm testing, the response to an unreinforced ambiguous arm (A) was tested. The full test sequence of the Test phase consisted therefore of 13 trials in total, with five P trials, five N trials, and three cue tests (e.g. P P N N P N **P** P N **N** P N **A**; cue testing in bold). The ambiguous arm (A) was spatially located midway between the two reference arms (N and P; 90°) and was associated with a mixed colored card (i.e. half green and red). Fish that failed to discriminate between P and N arms (as determined by showing a difference < 20 s in the latency between N and P) were retrospectively excluded from subsequent analyses.

Open Field and Novel Object Tests **-** Experimental fish were first exposed to a novel environment (Open Field Test; OFT), in an experimental apparatus consisting of a circular glass tank (21 cm diameter) with white walls in order to provide a consistent testing background and prevent access to external visual cues. The testing arena was filled with 2 L of water before the start of the experiment. Each fish was then gently placed in center of the tank and video recorded. Ten minutes later, a novel object (marble) was placed in the center of the tank to perform the Novel Object Test (NOT). Fish were then video recorded for additional 10 min.

Supplemental Note 5

Behavioral measures from OFT and NOT **-** During analysis, the OFT arena was virtually divided into an inner and an outer zone, with the outer zone defined as the area comprising one zebrafish body-length from the wall of the tank. For each fish, five behavioral measures were quantified: (1) distance moved (cm); (2) number of crossings between the inner and outer zones; (3) time spent in the inner zone (s); (4) distance from walls (cm); and (5) angular velocity (deg/s). The NOT arena was also virtually divided into two areas: the novel object and the empty zones, with the former defined as the area comprising one zebrafish body-length from the marble. Five behavioral measurements were also quantified: (1) distance moved (cm); (2) latency to first approach to the novel object (s); (3) number of approaches to the novel object; (4) time spent in the novel object zone (s); and (5) angular velocity (deg/s).

Ethics statement

All procedures were performed in accordance with the relevant guidelines and regulations for animal experimentation, reviewed by the Instituto Gulbenkian de Ciência Ethics Committee and by the institutional Organ Responsible for Animal Welfare (“Órgão Responsável pelo Bem-Estar dos Animais”, ORBEA) and approved by the competent Portuguese authority (Direcção Geral de Alimentação e Veterinária; permit number 0421/000/000/2019).

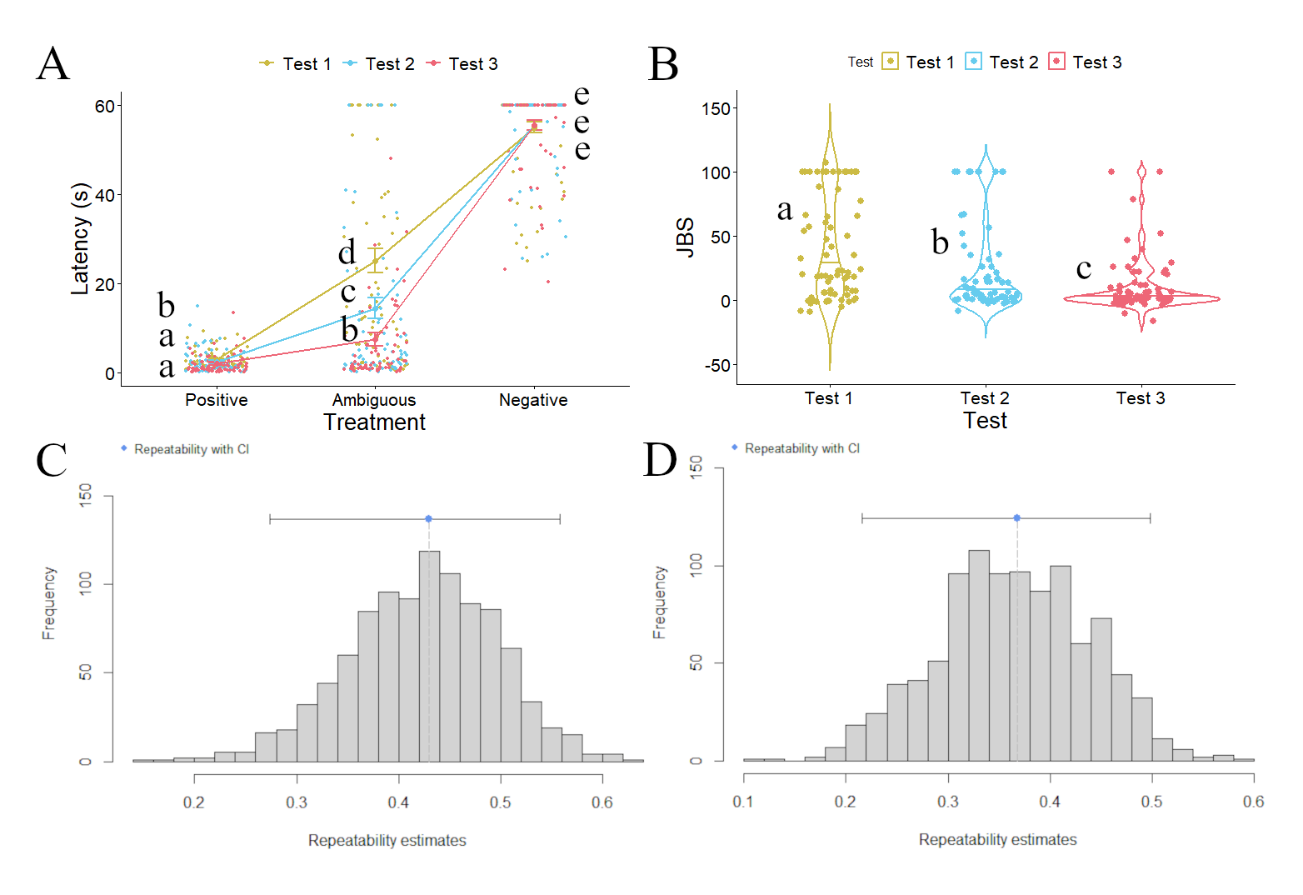
**Supplemental Figures and Tables**

Fig. S1. Behavioral characterization of judgment bias in zebrafish considering the whole population (n = 73). (A) Performance of male WT zebrafish in the judgement bias paradigm with repeated testing. Different letters indicate significant differences between experimental groups (Test 1, Test 2, and Test 3) for each Treatment (P, A, N) following planned comparisons tests. Data are expressed as mean ± s.e.m. (B) JBS of male WT zebrafish with repeated testing. Different letters indicate significant differences between experimental groups (Test 1, Test 2, and Test 3) following planned comparisons tests. Data are expressed as mean ± s.e.m. (C) Bootstrap repeatabilities for the latency to enter the ambiguous arm of the behavioral apparatus. (D) Bootstrap repeatabilities for the JBS.

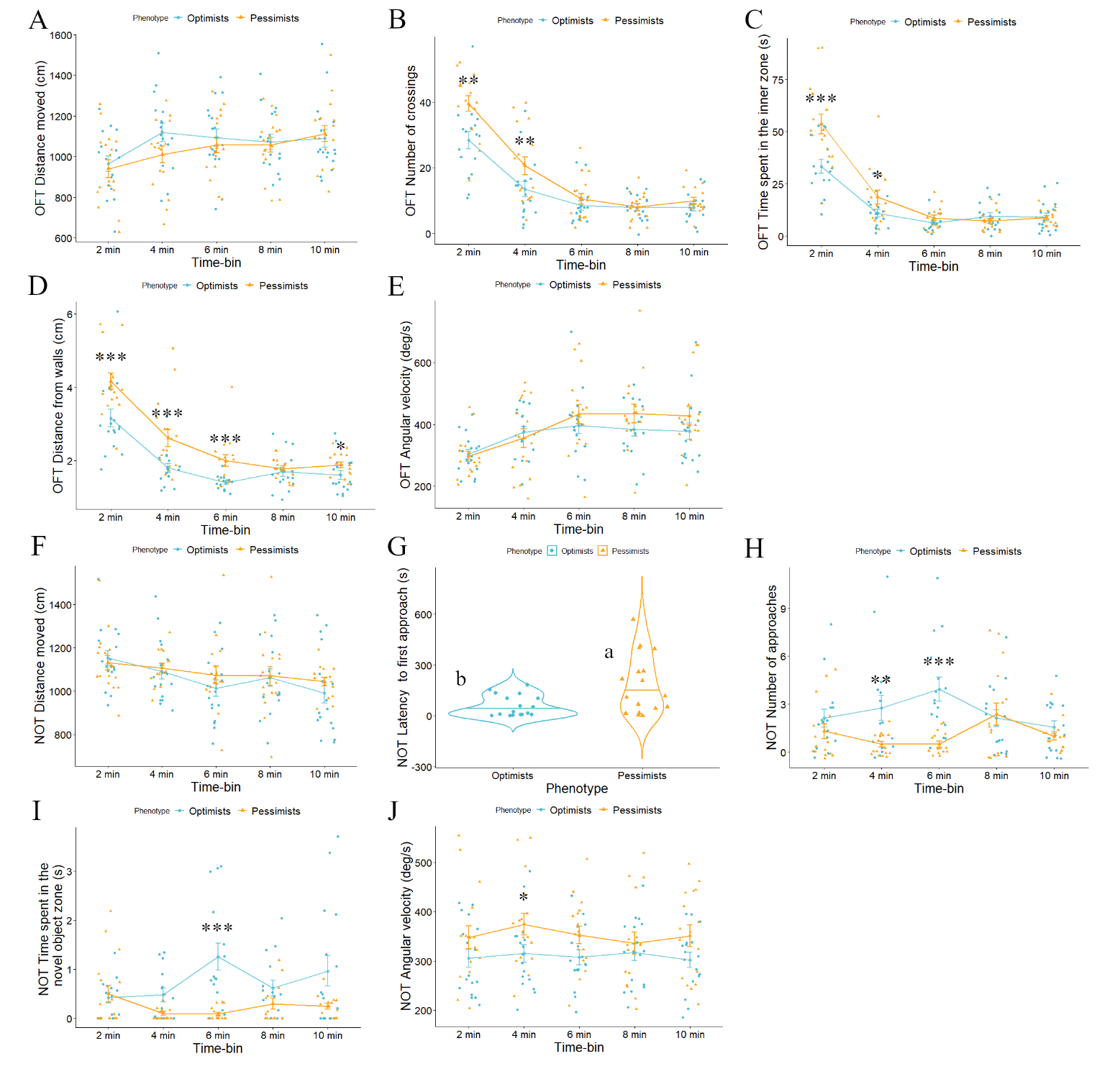
Fig. S2. Behavioral differences between fish from the upper and lower quartiles of the Judgment Bias Score (JBS; n = 17 per experimental group: optimists and pessimists): (A) distance moved (cm) in OFT; (B) number of crossings between the inner and outer zones in OFT; (C) time spent in the inner zone (s) in OFT; (D) distance from walls (cm) in OFT; (E) angular velocity (deg/s) in OFT; (F) distance moved (cm) in NOT; (G) latency to first approach to the novel object (s) in NOT; (H) number of approaches to the novel object in NOT; (I) time spent in the novel object zone (s) in NOT; (J) angular velocity (deg/s) in NOT. Asterisks and different letters indicate significant differences between the experimental groups at each sampling point following planned comparisons tests. Data are expressed as mean ± s.e.m.

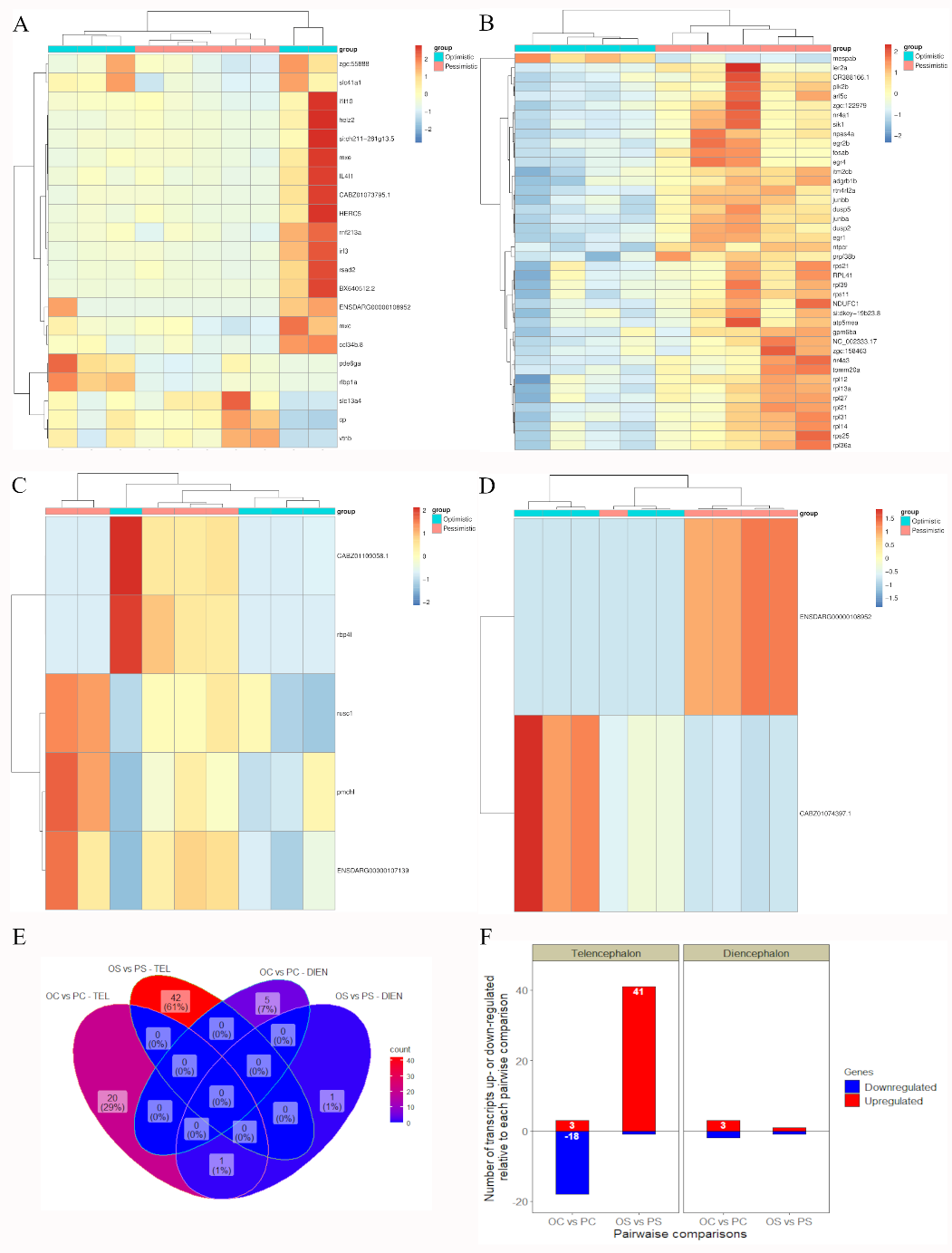

Fig. S3. Phenotypically driven changes in gene expression in the brain of control and stressed zebrafish (n = 5 per experimental group: optimists control, pessimists control, optimists stress, and pessimists stress). (A) Hierarchical clustering of control individuals from each phenotype (optimists versus pessimists; columns) and of DEG (lines) in the telencephalon; (B) Hierarchical clustering of stressed individuals from each phenotype (optimists versus pessimists; columns) and of DEG (lines) in the telencephalon; (C) Hierarchical clustering of control individuals from each phenotype (optimists versus pessimists; columns) and of DEG (lines) in the diencephalon; (D) Hierarchical clustering of pessimistic stressed from each phenotype (optimists versus pessimists; columns) and of DEG (lines) in the diencephalon. Heatmaps represent normalized gene expression levels (red, high expression; blue, low expression); (E) Venn diagram showing the number of DEG shared between each optimistic group and their pessimistic counterparts (optimists versus pessimists) in the telencephalon and diencephalon; (F) Total number of DEG up- and down-regulated relative to each optimistic group and their stressed counterparts (optimists versus pessimists) in the telencephalon and diencephalon.

**Table S1.** Results of the general linear mixed model to assess the effects of Test (Test 1 versus Test 2 versus Test 3), Treatment (Positive versus Ambiguous versus Negative), and the double interaction among these variables. Partial Eta Squared estimates of effect sizes are given for these factors. *Indicates a significant effect.

| Latency (s) upper and lower quartiles | | | | |
| --- | --- | --- | --- | --- |
| Main effects and interactions | F-value | p (>F) | Partial Eta^2^ | 95% CI |
| Test | F_2,264_=10.34 | p < 0.001* | 0.07 | [0.03, 1.00] |
| Treatment | F_2,264_=339.77 | p < 0.001* | 0.72 | [0.68, 1.00] |
| Test X Treatment | F_2,264_=5.18 | p < 0.001* | 0.07 | [0.02, 1.00] |
| Latency (s) whole population | | | | |
| Main effects and interactions | F-value | p (>F) | Partial Eta^2^ | 95% CI |
| Test | F_2,576_=38.25 | p < 0.001* | 0.12 | [0.08, 1.00] |
| Treatment | F_2,576_=965.84 | p < 0.001* | 0.77 | [0.75, 1.00] |
| Test Χ Treatment | F_2,576_=13.78 | p < 0.001* | 0.09 | [0.05, 1.00] |

**Table S2.** Results of the two-way ANOVA to assess the effects of Phenotype (Optimists versus Pessimists), Time (2 min versus 4 min versus 6 min versus 8 min versus 10 min), and the double interaction among these variables. Partial Eta Squared estimates of effect sizes are given for these factors. *Indicates a significant effect.

| OFT Distance moved (cm) | | | | |
| --- | --- | --- | --- | --- |
| Main effects and interactions | F-value | P-value | Partial Eta^2^ | 95% CI |
| Phenotype | F_1_ = 1.566 | p = 0.213 | 9.69e-03 | [0.00, 1.00] |
| Time | F_4_ = 4.072 | p = 0.004* | 0.09 | [0.02, 1.00] |
| Phenotype X Time | F_4_ = 0.701 | p = 0.593 | 0.02 | [0.00, 1.00] |
| OFT Number of crossings | | | | |
| Main effects and interactions | F-value | P-value | Partial Eta^2^ | 95% CI |
| Phenotype | F_1_ = 12.242 | p < 0.001* | 0.08 | [0.02, 1.00] |
| Time | F_4_ = 56.742 | p < 0.001* | 0.60 | [0.51, 1.00] |
| Phenotype X Time | F_4_ = 1.484 | p = 0.210 | 0.04 | [0.00, 1.00] |
| OFT Time spent in the inner zone (s) | | | | |
| Main effects and interactions | F-value | P-value | Partial Eta^2^ | 95% CI |
| Phenotype | F_1_ = 10.246 | p = 0.002* | 0.07 | [0.02, 1.00] |
| Time | F_4_ = 69.000 | p < 0.001* | 0.65 | [0.57, 1.00] |
| Phenotype X Time | F_4_ = 3.432 | p = 0.010* | 0.08 | [0.01, 1.00] |
| OFT Distance from walls (cm) | | | | |
| Main effects and interactions | F-value | P-value | Partial Eta^2^ | 95% CI |
| Phenotype | F_1_ = 37.594 | p < 0.001* | 0.19 | [0.11, 1.00] |
| Time | F_4_ = 52.705 | p < 0.001* | 0.58 | [0.49, 1.00] |
| Phenotype X Time | F_4_ = 1.598 | p = 0.178 | 0.04 | [0.00, 1.00] |
| OFT Angular velocity (deg/s) | | | | |
| Main effects and interactions | F-value | P-value | Partial Eta^2^ | 95% CI |
| Phenotype  Time  Phenotype X Time | F_1_ = 2.987  F_1_ = 6.239  F_1_ = 0.736 | p = 0.086  p < 0.001*  p = 0.569 | 8.97e-03  0.16  0.02 | [0.00, 1.00]  [0.07, 1.00]  [0.00, 1.00] |
| NOT Distance moved (cm) | | | | |
| Main effects and interactions | F-value | P-value | Partial Eta^2^ | 95% CI |
| Phenotype | F_1_ = 1.506 | p = 0.222 | 9.74e-03 | [0.00, 1.00] |
| Time | F_4_ = 3.834 | p = 0.005* | 0.09 | [0.02, 1.00] |
| Phenotype X Time | F_4_ = 0.569 | p = 0.685 | 0.01 | [0.00, 1.00] |
| NOT number of approaches | | | | |
| Main effects and interactions | F-value | P-value | Partial Eta^2^ | 95% CI |
| Phenotype | F_1_ = 18.572 | p < 0.001* | 0.11 | [0.04, 1.00] |
| Time | F_4_ = 0.826 | p = 0.510 | 0.02 | [0.00, 1.00] |
| Phenotype X Time | F_4_ = 3.576 | p = 0.008* | 0.09 | [0.01, 1.00] |
| NOT Time spent in the novel object zone (s) | | | | |
| Main effects and interactions | F-value | P-value | Partial Eta^2^ | 95% CI |
| Phenotype | F_1_ = 21.535 | p < 0.001* | 0.14 | [0.06, 1.00] |
| Time | F_4_ = 1.076 | p = 0.371 | 0.03 | [0.00, 1.00] |
| Phenotype X Time | F_4_ = 2.996 | p = 0.021* | 0.08 | [0.01, 1.00] |
| NOT Angular velocity (deg/s) | | | | |
| Main effects and interactions | F-value | P-value | Partial Eta^2^ | 95% CI |
| Phenotype | F_1_ = 12.769 | p < 0.001* | 0.07 | [0.02, 1.00] |
| Time | F_4_ = 0.338 | p = 0.852 | 8.37e-03 | [0.00, 1.00] |
| Phenotype X Time | F_4_ = 0.299 | p = 0.878 | 7.42e-03 | [0.00, 1.00] |

**Table S3.** List of DEG in the telencephalon.

|  | **Gene Symbol** | **Gene ID** | **logFC** | **P value** |
| --- | --- | --- | --- | --- |
| **Optimistic control vs Pessimistic control** | IL4I1; Cp; rnf213a; ifit10; rsad2; ENSDARG00000108952; BX640512,2; Mxc; CABZ01073795.1; helz2; irf3; pde6ga; si:ch211-281g13.5; rlbp1a; Mxe; zgc:55888; Vtnb; ccl34b.8; slc41a1; HERC5; slc13a4 | ENSDARG00000056804; ENSDARG00000010312; ENSDARG00000099465; ENSDARG00000007467; ENSDARG00000004952; ENSDARG00000108952; ENSDARG00000095112; ENSDARG00000024789; ENSDARG00000010729; ENSDARG00000016527; ENSDARG00000076251; ENSDARG00000056791; ENSDARG00000074052; ENSDARG00000012504; ENSDARG00000014427; ENSDARG00000016538; ENSDARG00000053831; ENSDARG00000093098; ENSDARG00000070214; ENSDARG00000075785; ENSDARG00000059053 | -2,40903553; 1,131914176; -1,19532024; -1,88463881; -1,78597027; -6,87979327; -3,72446705; -1,38392102; -1,96106196; -1,91341775; -2,10788061; -1,37630356; -1,87698639; -1,37195994; -2,04049276; -2,69538369; 0,936314083; -1,89545492; -0,99126826; -1,95067625; 1,265347259 | 2,66E-09; 7,06E-08; 1,17E-07; 1,88E-07; 6,76E-07; 6,52E-07; 1,42E-06; 1,72E-06; 2,00E-06; 4,27E-06; 9,20E-06; 1,36E-05; 1,57E-05; 2,75E-05; 2,76E-05; 2,71E-05; 2,25E-05; 2,56E-05; 4,28E-05; 4,44E-05; 4,78E-05 |
| **Optimistic control vs Optimistic stress** | Fosab; npas4a; Avp; rnf213a; rpl29; fads2; rps29; egr4; rpl39; rps21; klf11a; si:dkey-4p15.3; CR388166.1; rps28; si:dkeyp-27c8.2; pdk2a; atp5f1e; btg2; prpf38b; per1a; ier2a; pde6gb; RPL41; mid1ip1b; zgc:122979; fbxl3l; atp5mea; BX323458.1; gadd45ba; Mtbl; rcvrn3; atp5mf; rtn4rl2a | ENSDARG00000031683; ENSDARG00000055752; ENSDARG00000058567; ENSDARG00000099465; ENSDARG00000077717; ENSDARG00000019532; ENSDARG00000041232; ENSDARG00000077799; ENSDARG00000036316; ENSDARG00000025850; ENSDARG00000030844; ENSDARG00000037403; ENSDARG00000087753; ENSDARG00000035860; ENSDARG00000088040; ENSDARG00000020876; ENSDARG00000095897; ENSDARG00000020298; ENSDARG00000053101; ENSDARG00000056885; ENSDARG00000099195; ENSDARG00000101984; ENSDARG00000092807; ENSDARG00000019302; ENSDARG00000004187; ENSDARG00000037738; ENSDARG00000078113; ENSDARG00000078220; ENSDARG00000027744; ENSDARG00000102051; ENSDARG00000009637; ENSDARG00000037867; ENSDARG00000052012 | -1,97774843; -1,95398998; 0,9310121; -1,16171969; -0,6878281; -0,59704996; -0,66665029; -1,32295521; -0,56804191; -0,59969524; -0,67077816; -0,65234135; -1,09411341; -0,57205725; -0,90636866; 0,735238704; -0,57432421; -1,0110221; -0,8500379; 1,170480423; -1,17963758; 1,658054979; -0,48645439; -0,61262808; -1,55150835; 1,102229001; -0,58901221; -1,85066318; -0,58067767; -0,55326269; 1,10038794; -0,50720642; -0,71851865 | 1,83E-11; 1,61E-09; 3,53E-09; 2,11E-08; 7,18E-07; 2,60E-06; 3,22E-06; 3,07E-06; 5,03E-06; 6,77E-06; 7,47E-06; 7,89E-06; 6,18E-06; 9,46E-06; 9,88E-06; 1,13E-05; 1,35E-05; 1,82E-05; 1,69E-05; 1,74E-05; 1,88E-05; 2,45E-05; 2,58E-05; 3,24E-05; 4,33E-05; 4,29E-05; 4,34E-05; 4,31E-05; 4,60E-05; 5,84E-05; 6,65E-05; 6,49E-05; 6,69E-05 |
| **Pessimistic control vs Pessimistic stress** | wu:fj16a03; c4; si:ch1073-266p11.2; cldn11a; opn1lw1; fxyd1; gpt2l; Ntpcr; kcnj2a; grk1b; krt8; slc20a1b; Smox; Apoeb; si:dkey-56d12.4; clec11a; Glulb; unc119,2; NCKAP1L; si:ch73-364h19.2; pde6ha; rcvrn2; nr1d4b; cpxm1a; rd3; Pltp; lamc3; CABZ01074397.1; Spra; opn1lw2; slc38a4; Pdcb; Pgr; iqgap1; pld6; Cp; ENSDARG00000107716; Bhmt; sulf1; si:dkeyp-106c3.1; slc22a7b.1; Lipg; nr1d4a; itga8; si:ch211-250k18.7; exd1; gngt2a; hoxa2b | ENSDARG00000100952; ENSDARG00000015065; ENSDARG00000100261; ENSDARG00000020031; ENSDARG00000044862; ENSDARG00000099014; ENSDARG00000019541; ENSDARG00000021853; ENSDARG00000019418; ENSDARG00000104685; ENSDARG00000058358; ENSDARG00000010641; ENSDARG00000036967; ENSDARG00000040295; ENSDARG00000070845; ENSDARG00000079107; ENSDARG00000100003; ENSDARG00000004459; ENSDARG00000075748; ENSDARG00000100480; ENSDARG00000102558; ENSDARG00000019902; ENSDARG00000059370; ENSDARG00000073716; ENSDARG00000031600; ENSDARG00000104495; ENSDARG00000093572; ENSDARG00000104672; ENSDARG00000004406; ENSDARG00000044861; ENSDARG00000018149; ENSDARG00000017634; ENSDARG00000035966; ENSDARG00000078888; ENSDARG00000059951; ENSDARG00000010312; ENSDARG00000107716; ENSDARG00000013430; ENSDARG00000038428; ENSDARG00000090623; ENSDARG00000056643; ENSDARG00000031044; ENSDARG00000031161; ENSDARG00000078717; ENSDARG00000093761; ENSDARG00000098669; ENSDARG00000010680; ENSDARG00000023031 | -1,28854371; -1,371538; 1,20095094; -1,79238955; 2,62811292; -1,36293034; -1,16877436; 1,20045282; -1,67395686; 2,04568767; -0,60913129; -0,49727238; -0,62662127; -1,06561755; -3,04133173; -1,71541424; -0,70312345; 2,11506302; -0,98373825; -8,22785989; 1,1095601; 1,37421725; 1,40864684; -1,44813561; 1,43368691; -1,02589239; -1,11579351; -2,87285259; 0,91078389; 1,58390578; -0,78281594; 1,74168941; -1,03957012; -0,67206348; 1,40566129; -1,95636626; 1,53909251; -1,39265027; -0,57928601; -1,556006; -1,39635318; -0,67141624; 0,68546359; -0,98955244; 1,84429895; -1,62539428; 1,15414243; 1,91472032 | 1,64E-11; 2,57E-10; 3,89E-09; 1,18E-08; 1,47E-08; 3,72E-08; 2,13E-07; 2,22E-07; 2,57E-07; 3,29E-07; 5,34E-07; 1,16E-06; 1,10E-06; 1,17E-06; 8,37E-07; 9,81E-07; 1,11E-06; 1,54E-06; 1,66E-06; 1,81E-06; 2,35E-06; 6,42E-06; 6,61E-06; 9,69E-06; 1,07E-05; 1,19E-05; 1,33E-05; 1,34E-05; 1,82E-05; 2,08E-05; 2,51E-05; 2,72E-05; 2,67E-05; 3,41E-05; 3,59E-05; 3,97E-05; 4,08E-05; 4,67E-05; 4,79E-05; 4,87E-05; 5,00E-05; 5,38E-05; 6,40E-05; 6,81E-05; 6,81E-05; 7,45E-05; 9,02E-05; 0,00010843 |
| **Optimistic stress vs Pessimistic stress** | rtn4rl2a; nr4a1; plk2b; dusp5; Junba; dusp2; itm2cb; egr1; CR388166.1; Fosab; NC_002333.17; Junbb; arl5c; nr4a3; npas4a; Ntpcr; rpl39; adgrb1b; rps25; egr2b; ier2a; rpl21; rps11; Mespab; egr4; si:dkey-19b23.8; rpl12; rpl36a; prpf38b; NDUFC1; atp5mea; rpl31; zgc:122979; rpl13a; rpl14; sik1; tomm20a; rps21; zgc:158463; RPL41; gpm6ba; rpl27 | ENSDARG00000052012; ENSDARG00000000796; ENSDARG00000019130; ENSDARG00000019307; ENSDARG00000074378; ENSDARG00000098108; ENSDARG00000039650; ENSDARG00000037421; ENSDARG00000087753; ENSDARG00000031683; ENSDARG00000082753; ENSDARG00000104773; ENSDARG00000035719; ENSDARG00000055854; ENSDARG00000055752; ENSDARG00000021853; ENSDARG00000036316; ENSDARG00000078529; ENSDARG00000041811; ENSDARG00000042826; ENSDARG00000099195; ENSDARG00000010516; ENSDARG00000053058; ENSDARG00000068761; ENSDARG00000077799; ENSDARG00000103211; ENSDARG00000006691; ENSDARG00000058105; ENSDARG00000053101; ENSDARG00000103101; ENSDARG00000078113; ENSDARG00000053365; ENSDARG00000004187; ENSDARG00000044093; ENSDARG00000103433; ENSDARG00000058606; ENSDARG00000090656; ENSDARG00000025850; ENSDARG00000089382; ENSDARG00000092807; ENSDARG00000005739; ENSDARG00000015128 | 0,92920828; 1,94297902; 0,88926321; 0,89530165; 0,91781152; 0,95334498; 0,71586851; 0,68694999; 1,15478946; 2,25890604; 0,66435257; 0,66900708; 1,00869068; 0,8527366; 1,85497038; 1,07123856; 0,52843009; 0,50626335; 0,54793254; 1,79540868; 1,58006574; 0,45697247; 0,47833896; -2,0854341; 1,46201332; 0,67096888; 0,49149804; 0,50029147; 0,68971041; 0,52119153; 0,53868348; 0,43100925; 1,73485108; 0,40649445; 0,47351797; 0,5779233; 1,08732136; 0,48694527; 0,57091796; 0,41588808; 0,55875563; 0,43227459 | 2,40E-13; 5,79E-12; 1,69E-11; 1,18E-11; 1,76E-11; 2,62E-10; 9,10E-09; 1,10E-08; 4,79E-08; 6,33E-08; 7,47E-08; 2,17E-07; 2,68E-07; 3,25E-07; 8,36E-07; 3,98E-06; 5,00E-06; 6,32E-06; 7,09E-06; 7,62E-06; 8,80E-06; 1,64E-05; 1,63E-05; 1,66E-05; 1,62E-05; 1,96E-05; 2,75E-05; 3,21E-05; 3,41E-05; 3,93E-05; 4,55E-05; 4,82E-05; 5,07E-05; 5,62E-05; 5,53E-05; 6,36E-05; 7,17E-05; 7,85E-05; 8,25E-05; 8,34E-05; 8,67E-05; 9,82E-05 |

**Table S4.** List of DEG in the diencephalon.

|  | **Gene Symbol** | **Gene ID** | **logFC** | **P value** |
| --- | --- | --- | --- | --- |
| **Optimistic control vs Pessimistic control** | CABZ01109058.1; Pmchl; rbp4l; ENSDARG00000107139; rusc1 | ENSDARG00000103574; ENSDARG00000076978; ENSDARG00000044684; ENSDARG00000107139; ENSDARG00000078125 | -1,10329742; 1,50461433; -1,06036437; 2,00997031; 0,60652528 | 1,66E-08; 2,28E-07; 1,94E-06; 3,47E-06; 7,43E-06 |
| **Optimistic control vs Optimistic stress** | kctd3; hsp90aa1.2; si:ch211-117k10.3; sept4a; Pmchl; tlcd1; prlh2; msmo1; CR936442.1; fa2h; CABZ01074397.1; cyp51; zgc:92630; ENSDARG00000069159; bcl6b; pycr1a; sptlc2b; BX936284.1; cldn19; per1a; si:dkey-170l10.1; si:dkey-145p14.5; klf11a; mid1ip1b; sc5d; prss35; mcm7; plxna1a; pth2; si:ch211-132b12.7; ppp1r14aa; si:ch211-237l4.6; Sqlea; si:ch211-132g1.6; cdkn1d; ENSDARG00000034092; tubb5; Hells; hsd17b7; scarb2a; plp1b; ENSDARG00000108666; clk4a; tecpr2; Mpz; rbp4l; tpx2; zgc:63568; si:dkey-56m19.5; eepd1 | ENSDARG00000060854; ENSDARG00000024746; ENSDARG00000090914; ENSDARG00000105271; ENSDARG00000076978; ENSDARG00000002391; ENSDARG00000071735; ENSDARG00000055876; ENSDARG00000105592; ENSDARG00000090063; ENSDARG00000104672; ENSDARG00000042641; ENSDARG00000004141; ENSDARG00000069159; ENSDARG00000069335; ENSDARG00000102254; ENSDARG00000074287; ENSDARG00000101826; ENSDARG00000044569; ENSDARG00000056885; ENSDARG00000036781; ENSDARG00000007077; ENSDARG00000030844; ENSDARG00000019302; ENSDARG00000044642; ENSDARG00000100691; ENSDARG00000101180; ENSDARG00000105452; ENSDARG00000022951; ENSDARG00000068374; ENSDARG00000011239; ENSDARG00000056915; ENSDARG00000079946; ENSDARG00000093102; ENSDARG00000099719; ENSDARG00000034092; ENSDARG00000037997; ENSDARG00000057738; ENSDARG00000088140; ENSDARG00000098312; ENSDARG00000011929; ENSDARG00000108666; ENSDARG00000089372; ENSDARG00000060835; ENSDARG00000038609; ENSDARG00000044684; ENSDARG00000078654; ENSDARG00000087262; ENSDARG00000068432; ENSDARG00000071116 | 1,6439833; -0,80377713; 1,03980043; -0,69766415; 1,70469478; -0,81239356; 1,29731506; -0,5667235; -0,81471085; -0,57272972; 2,50542942; -0,63500416; -0,76087655; 1,60236252; -0,71042438; 0,84429321; -0,63713926; -0,50661076; -0,45873325; 0,98474479; -1,07463863; -0,51107445; -0,56966392; -0,59512731; -0,51448116; -0,66977549; -0,9819836; 0,5257836; 0,70015012; -0,70841371; -0,51714799; -0,58303156; -0,65168711; 1,66364808; 0,73977834; 1,39589095; -0,4320494; -0,81381546; -0,48070541; -0,58709474; -0,51370133; -0,59034314; -0,42122872; 0,5004126; -0,41499193; -0,82396956; -0,95280375; -1,23256411; -0,44611144; -0,61524309 | 3,98E-15; 2,15E-09; 3,55E-09; 1,02E-07; 1,41E-07; 3,13E-07; 2,74E-07; 7,25E-07; 1,32E-06; 1,60E-06; 2,61E-06; 4,34E-06; 4,94E-06; 7,52E-06; 7,27E-06; 7,48E-06; 8,51E-06; 8,90E-06; 1,03E-05; 1,07E-05; 1,21E-05; 1,76E-05; 1,79E-05; 2,29E-05; 2,01E-05; 2,12E-05; 2,24E-05; 2,15E-05; 3,05E-05; 3,22E-05; 3,49E-05; 3,75E-05; 3,82E-05; 4,12E-05; 4,04E-05; 4,30E-05; 5,11E-05; 5,07E-05; 5,19E-05; 5,30E-05; 5,44E-05; 5,60E-05; 6,26E-05; 7,05E-05; 8,51E-05; 8,59E-05; 8,35E-05; 8,33E-05; 0,00011388; 0,00011394 |
| **Pessimistic control vs Pessimistic stress** | nr1d4b; ENSDARG00000108952; fbxl3l; si:dkey-56d12.4; ENSDARG00000075877; gpt2l; snu13a; si:ch1073-266p11.2; CABZ01109058.1; pde6a | ENSDARG00000059370; ENSDARG00000108952; ENSDARG00000037738; ENSDARG00000070845; ENSDARG00000075877; ENSDARG00000019541; ENSDARG00000069878; ENSDARG00000100261; ENSDARG00000103574; ENSDARG00000000380 | 2,85144891; 5,73038448; 1,12879117; -3,26162779; 1,52695987; -0,60001119; -0,64487506; 1,18916912; 0,90093213; 0,84011205 | 6,18E-09; 1,47E-08; 6,04E-07; 7,17E-07; 1,13E-06; 3,94E-06; 7,89E-06; 7,87E-06; 8,18E-06; 1,09E-05 |
| **Optimistic stress vs Pessimistic stress** | ENSDARG00000108952; CABZ01074397.1 | ENSDARG00000108952; ENSDARG00000104672 | 5,85471713; -2,93139926 | 5,43E-10; 5,29E-09 |

|  | **GOBP** | **GOCC** | **GOMF** |
| --- | --- | --- | --- |
| **Optimistic control vs Pessimistic control** | Multi-organism process (22%)  Mitochondrial fission (11%)  Response to stimulus (39%)  Dynamin family protein polymerization involved in mitochondrial fission (11%)  Immune system process (33%)  Defense response (28%)  Immune response (28%)  Defense response to virus (22%) | - | - |
| **Optimistic control vs Optimistic stress** | Translation (25%) | Ribosome (32%)  Rough endoplasmic reticulum (11%)  Cytosolic part (26%)  Ribosomal subunit (26%) | Structural molecule activity (25%)  Structural constituent of ribosome (25%) |
| **Pessimistic control vs Pessimistic stress** | - | - | - |
| **Optimistic stress vs Pessimistic stress** | Cellular response to a hormone stimulus (6.3%)  Negative regulation of intracellular signal transduction (9.4%)  Metabolic process (78%)  Formation of primary germ layer (9.4%)  Lymph vessel development (6.3%)  Biosynthetic process (63%)  Regulation of macromolecule metabolic process (41%)  Organic substance biosynthetic process (63%)  Cellular biosynthetic process (63%)  Negative regulation of protein phosphorylation (9.4%)  Macromolecule biosynthetic process (59%)  Cellular nitrogen compound biosynthetic process (63%)  Negative regulation of protein kinase activity (9.4%)  Transcription, DNA-template (25%)  Translation (34%) | Protein-containing complex (52%)  Organelle (68%)  Intracellular (29%)  Ribosome (39%)  Organelle part (42%)  Rough endoplasmic reticulum (6.5%)  Ribosomal subunit (32%)  Cytosolic large ribosomal subunit (26%) | Nucleic acid binding (44%)  Transcription regulation activity (21%)  MAO kinase tyrosine/serine/threonine phosphatase activity (5.9%)  Structural molecule activity (32%)  RNA polymerase II proximal promoter sequence-specific DNA binding (8.8%)  DNA binding transcription factor activity (21%)  Structural constituent of ribosome (32%) |

**Table S5.** Characterization of the DE genes in the telencephalon obtained using ORA for GO, and summarized using GOSlim terms.

**Table S6.** Characterization of the DE genes in the diencephalon obtained using ORA for GO, and summarized using GOSlim terms.

|  | **GOBP** | **GOCC** | **GOMF** |
| --- | --- | --- | --- |
| **Optimistic control vs Pessimistic control** | - | - | - |
| **Optimistic control vs Optimistic stress** | Sterol biosynthetic process (9.1%)  Secondary alcohol metabolic process (9.1%)  Response to antibiotic (6.1%)  Response to yeast (6.1%)  Cholesterol metabolic process (9.1%) | - | - |
| **Pessimistic control vs Pessimistic stress** | - | - | - |
| **Optimistic stress vs Pessimistic stress** | - | - | - |

**Table S7.** Results of the general linear mixed model to assess the effects of Phenotype (Optimists versus Pessimists), Treatment (Control versus Stress), and the double interaction among these variables. Partial Eta Squared estimates of effect sizes are given for these factors. *Indicates a significant effect.

| Whole-body cortisol (ng/g) | | | | |
| --- | --- | --- | --- | --- |
| Main effects and interactions | F-value | p (>F) | Partial Eta^2^ | 95% CI |
| Phenotype | F_1,33.9_ = 1.638 | p = 0.209 | 0.05 | [0.00, 1.00] |
| Treatment | F_1,6.9_ = 5.115 | p = 0.058 | 0.42 | [0.00, 1.00] |
| Phenotype X Treatment | F_1,33.9_ = 0.822 | p = 0.370 | 0.02 | [0.00, 1.00] |
| *crh* Telencephalon | | | | |
| Main effects and interactions | F-value | p (>F) | Partial Eta^2^ | 95% CI |
| Phenotype | F_1,34.7_ = 0.032 | p = 0.858 | 9.21e-04 | [0.00, 1.00] |
| Treatment | F_1,7.4_ = 5.309 | p = 0.052 | 0.41 | [0.00, 1.00] |
| Phenotype X Treatment | F_1,34.7_ = 0.030 | p = 0.861 | 8.87e-04 | [0.00, 1.00] |
| *gr* Telencephalon | | | | |
| Main effects and interactions | F-value | p (>F) | Partial Eta^2^ | 95% CI |
| Phenotype | F_1,39_ = 0.885 | p = 0.352 | 0.02 | [0.00, 1.00] |
| Treatment | F_1,39_ = 0.804 | p = 0.375 | 0.02 | [0.00, 1.00] |
| Phenotype X Treatment | F_1,39_ = 7.561 | p = 0.008* | 0.16 | [0.03, 1.00] |
| *mr* Telencephalon | | | | |
| Main effects and interactions | F-value | p (>F) | Partial Eta^2^ | 95% CI |
| Phenotype | F_1,39_ = 5.081 | p = 0.029* | 0.12 | [0.01, 1.00] |
| Treatment | F_1,39_ = 0.121 | p = 0.729 | 3.10e-03 | [0.00, 1.00] |
| Phenotype X Treatment | F_1,39_ = 8.480 | p = 0.005* | 0.18 | [0.03, 1.00] |
| *mr/gr* Telencephalon | | | | |
| Main effects and interactions | F-value | p (>F) | Partial Eta^2^ | 95% CI |
| Phenotype  Treatment  Phenotype X Treatment | F_1,32.3_ = 10.294  F_1,7.7_ = 3.703  F_1,32.2_ = 1.729 | p = 0.003*  p = 0.091  p = 0.197 | 0.24  0.32  0,05 | [0.06, 1.00]  [0.00, 1.00]  [0.00, 1.00] |
| *crh* Diencephalon | | | | |
| Main effects and interactions | F-value | p (>F) | Partial Eta^2^ | 95% CI |
| Phenotype | F_1,41_ = 0.022 | p = 0.882 | 5.43e-04 | [0.00, 1.00] |
| Treatment | F_1,41_ = 6.493 | p = 0.014* | 0.14 | [0.02, 1.00] |
| Phenotype X Treatment | F_1,41_ = 0.144 | p = 0.705 | 3.51e-03 | [0.00, 1.00] |
| *gr* Diencephalon | | | | |
| Main effects and interactions | F-value | p (>F) | Partial Eta^2^ | 95% CI |
| Phenotype | F_1,41_ = 5.099 | p = 0.029* | 0.11 | [0.01, 1.00] |
| Treatment | F_1,41_ = 6.729 | p = 0.013* | 0.14 | [0.02, 1.00] |
| Phenotype X Treatment | F_1,41_ = 2.099 | p = 0.154 | 0.05 | [0.00, 1.00] |
| *mr* Diencephalon | | | | |
| Main effects and interactions | F-value | p (>F) | Partial Eta^2^ | 95% CI |
| Phenotype | F_1,41_ = 0.737 | p = 0.395 | 0.02 | [0.00, 1.00] |
| Treatment | F_1,41_ = 0.157 | p = 0.693 | 3.83e-03 | [0.00, 1.00] |
| Phenotype X Treatment | F_1,41_ = 2.222 | p = 0.143 | 0.05 | [0.00, 1.00] |
| *mr/gr* Diencephalon | | | | |
| Main effects and interactions | F-value | p (>F) | Partial Eta^2^ | 95% CI |
| Phenotype | F_1,41_ = 15.411 | p < 0.001* | 0.27 | [0.10, 1.00] |
| Treatment | F_1,41_ = 3.116 | p = 0.084 | 0.07 | [0.00, 1.00] |
| Phenotype X Treatment | F_1,41_ = 1.162 | p = 0.287 | 0.03 | [0.00, 1.00] |

**Table S8.** Results of the general linear mixed model to assess the effects of Genotype (mitfa:HRAS^G12V^-GFP versus WT), Treatment (Positive versus Ambiguous versus Negative), and the double interaction among these variables. Partial Eta Squared estimates of effect sizes are given for these factors. *Indicates a significant effect.

| Latency (s) | | | | |
| --- | --- | --- | --- | --- |
| Main effects and interactions | F-value | p (>F) | Partial Eta^2^ | 95% CI |
| Genotype | F_1,155_=28.09 | p < 0.001* | 0.15 | [0.08, 1.00] |
| Treatment | F_2,155_=112.76 | p < 0.001* | 0.59 | [0.51, 1.00] |
| Genotype X Treatment | F_2,155_=6.97 | p = 0.0012* | 0.08 | [0.02, 1.00] |

**Table S9**. Comparison of the survival curves of the different experimental groups. Treatments were compared using the log-rank test. Correction of p values was performed using the Holm-Sidak method. *Indicates a significant effect.

| Treat 1 | Treat 2 | p-value | Adjusted p-value |
| --- | --- | --- | --- |
| OC | PC | 0.011676 | 0.045892* |
| OC | OS | 0.000001 | 0.000008* |
| PC | PS | 0.080082 | 0.153751 |
| OS | PS | 0.384289 | 0.384289 |

**Table S10.** Comparison of the survival probability at specific timepoints between treatments that were significantly different**.** Note that when the survival curve is zero the statistic is infinite. *Indicates a significant effect.

| Comparison | Time point | Statistic | p-value | Adjusted p-value |
| --- | --- | --- | --- | --- |
| OC – PC | 9 weeks | 5.894953 | 0.015184 | 0.043613* |
| OC – PC | 10 weeks | 5.945593 | 0.014754 | 0.043613* |
| OC – PC | 11 weeks | 1.130744 | 0.287617 | 0.287617 |
| OC – OS | 9 weeks | 11.788144 | 0.000596 | 0.002977* |
| OC – OS | 10 weeks | infinite | 0.000000 | 0.000000* |
| OC – OS | 11 weeks | infinite | 0.000000 | 0.000000* |

| Tumor fraction (%) | | | | |
| --- | --- | --- | --- | --- |
| Main effects and interactions | F-value | p (>F) | Partial Eta^2^ | 95% CI |
| Phenotype | F_1,10.38_ = 0.197 | p = 0.665 | 0.02 | [0.00, 1.00] |
| Treatment | F_1,10.38_ = 5.881 | p = 0.034* | 0.36 | [0.02, 1.00] |
| Phenotype X Treatment | F_1,10.38_= 1.033 | p = 0.332 | 0.09 | [0.00, 1.00] |
| PCNA positive tumor cells (%) | | | | |
| Main effects and interactions | F-value | p (>F) | Partial Eta^2^ | 95% CI |
| Phenotype | F_1,10.24_ = 10.155 | p = 0.009* | 0.50 | [0.11, 1.00] |
| Treatment | F_1,10.24_ = 0.014 | p = 0.905 | 1.45e-03 | [0.00, 1.00] |
| Phenotype X Treatment | F_1,10.24_ = 1.069 | p = 0.324 | 0.09 | [0.00, 1.00] |

**Table S11.** Results of the general linear mixed model to assess the effects of Phenotype (Optimists versus Pessimists), Treatment (Control versus Stress), and the interaction among these variables**.** Partial Eta Squared estimates of effect sizes are given for these factors. *Indicates a significant effect.

| Table S12. Procedure of the unpredictable chronic stress (UCS) protocol in zebrafish. | | | | | | |
| --- | --- | --- | --- | --- | --- | --- |
| Day 1 | **Day 2** | **Day 3** | **Day 4** | **Day 5** | **Day 6** | **Day 7** |
| 9:00h Overcrowding  12:00h Heat up water | 14:00h Social Isolation  17:30h Alarm substance | 16:00h Restraint stress  18:00h Cool down water | 9:00h Air exposure  17:30h Tank changes | 10:30h Social Isolation  15:00h Overcrowding | 10:00h Heat up water  16:30h Alarm substance | 12:00h Restraint stress  17:30h Overcrowding |
| Day 8 | **Day 9** | **Day 10** | **Day 11** | **Day 12** | **Day 13** | **Day 14** |
| 12:30h Air exposure  18:30h Chasing | 14:00h Min water level  16:30h Tank changes | 11:30h Cool down water  17:30h Min water level | 10:00h Heat up water  18:30h Chasing | 16:30h Tank changes  17:30h Alarm substance | 9:00h Restraint stress  11:30h Overcrowding | 14:30h Min water level  17:00h Air exposure |
| Day 15 | **Day 16** | **Day 17** | **Day 18** | **Day 19** | **Day 20** | **Day 21** |
| 9:00h Social Isolation  18:30h Chasing | 12:00h Heat up water  17:00h Cool down water | 9:00h Tank changes  11:30h Cool down water | 17:00h Overcrowding  18:00h Chasing | 11:00h Min water level  16:30h Restraint stress | 12:00h Tank changes  15:30h Alarm substance | 14:00h Heat up water  18:30h Air exposure |
| Day 22 | **Day 23** | **Day 24** | **Day 25** | **Day 26** | **Day 27** | **Day 28** |
| 9:30h Social Isolation  16:00h Chasing | 11:30h Social Isolation  13:30h Restraint stress | 12:00h Cool down water  17:00h Alarm substance | 15:00h Chasing  16:30h Cool down water | 13:00h Min water level  18:30h Tank changes | 17:00h Min water level  18:30h Overcrowding | 9:00h Social Isolation  16:30h Alarm substance |
| Day 29 | **Day 30** |  |  |  |  |  |
| 14:30h Heat up water  16:00h Air exposure | 10:00h Air exposure  17:00h Restraint stress |  |  |  |  |  |

**Table S13.** Primer sequences ^a^, amplicon lengths and annealing parameters for the genes used in the qPCR.

| Gene | NCBI Reference Sequence | Primer sequence (5’→3’) | Apmlicon size (pb) | Temperature of annealing (°C) | Time of annealing (s) | PCR efficiency |
| --- | --- | --- | --- | --- | --- | --- |
| *ef1-alpha* | NM_131263.1 | ^b^ fw: CAAGGAAGTCAGCGCATACA  ^c^ rv: TCTTCCATCCCTTGAACCAG | 134 | 59 | 30 | 0.91 |
| *Crh* | NM_001007379.1 | fw: GGCAACAGAAACCCGACTT  rv: CAACTTTCCCCTCCAACAGA | 118 | 61 | 30 | 0.98 |
| *Gr* | NM_001020711.3 | fw: GCTCAATGGCACAGCTTCTT  rv: CCGGTGTTCTCCTGTTTGAT | 126 | 59 | 30 | 1.03 |
| *Mr* | NM_001100403.1 | fw: CAACAACCGCAAGTCAGAAA  rv: TGTTGGGAAAAGCCAAAGTC | 111 | 62 | 60 | 0.96 |

^a^ Forward and reverse primers were obtained from Sigma-Aldrich (Hamburg, Germany).

^b^ fw: forward primer.

^c^ rv: reverse primer.
